# Phage Mediate Bacterial Self Recognition

**DOI:** 10.1101/413146

**Authors:** Sooyeon Song, Yunxue Guo, Jun-Seob Kim, Xiaoxue Wang, Thomas K. Wood

## Abstract

Cells are social, and self-recognition is an important and conserved aspect of group behavior where cells assist kin and antagonize non-kin to conduct group behavior such as foraging for food and biofilm formation. However, the role of the common bacterial cohabitant, phage, in kin recognition, has not been explored. Here we find that a boundary (demarcation line) is formed between different swimming *Escherichia coli* strains but not between identical clones; hence, motile bacterial cells discriminate between self and non-self. The basis for this self-recognition is a novel, 49 kb, T1-type, lytic phage of the family siphoviridae (named here SW1) that controls formation of the demarcation line by utilizing one of the host’s cryptic prophage proteins, YfdM, to propagate. Critically, SW1 increases the fitness of *E. coli* K-12 compared to the identical strain that lacks the phage. Therefore, bacteria use phage to recognize kin.

---

Cellular motility impacts bacterial physiology by providing a means for acquiring nutrients (e.g., chemotaxis)^1^, evading stress^1^, initiating biofilm formation^2^, and dispersing biofilms^3^. Two of the best-studied forms of motility are swarming, which is flagella-based movement *on top* of a solid surface, and swimming, which is flagella-based movement *through* liquids^4^. Motility can also be social and is a critical part of self-recognition, a behavior that impacts nourishment, virulence, iron acquisition, protection, quorum sensing, biofilm formation, and fruiting bodies^5^. In effect, self-recognition allows bacteria to form social groups^6^.

Self-recognition mechanisms studied to date depend on surface receptors and diffusible chemical signals^7^. For example, during the swarming of *Proteus mirabilis*, self-recognition takes the form of a demarcation line (Dienes line^8^); as cells move on the surface, a visible gap is formed between different strains due to killing as a result of cell-cell contact through identity proteins (IdsD and IdsE)^9^ as well as due to killing by secreted bacteriocins^5^. Similarly, several self-recognition determinants have been identified for swarming *Bacillus subtilis* cells including contact-dependent inhibition and secreted antimicrobials^10^.

Additionally, self-recognition during swarming is mediated through a toxin-antitoxin system requiring outer membrane exchange in *Myxococcus xanthus*, in which non-motile cells can prevent the swarming of cells that lack the antitoxin^11^. However, self-recognition during swimming motility (i.e., motility *through* liquids) has not been reported previously, and self-recognition during swimming or swarming has not been linked to phage-mediated cell lysis.

*Escherichia coli* K-12 contains nine cryptic prophages, that individually are incapable of producing active phage particles, and constitute 3.6% of its genome^12^. Rather than genomic junk, we found these cryptic prophages help the cell cope with stress^12^. We also found that prophage genes are induced in *E. coli* during biofilm formation^13^ and that cryptic prophage influence biofilm formation in this strain^12, 14, 15^. Furthermore, active, filamentous phage influence the biofilm dispersal and phenotypic variation of *Pseudomonas aeruginosa*^16^. Hence, cryptic and active phages are involved intimately in cell physiology in roles distinct from phage propagation.

Here, we became intrigued when we encountered a demarcation line between different swimming *E. coli* cells (**Fig. 1**) and investigated the underlying mechanism. We screened the swimming behavior of 4296 single-gene knockouts of the Keio Collection^17^ and found that the demarcation line disappears for a mutation related to phage development. By hypothesizing the gap between swimming cells was related to phage lysis, we identified a novel T1-type lytic phage (named SW1) that allows *E. coli* K-12 to recognize kin (this phage appears to arise from the Keio Collection). By utilizing our strains devoid of each of the nine resident cryptic prophage in *E. coli*^12^, we determined that the demarcation line is enhanced by putative methylase YfdM of cryptic phage CPS-53, and the extent of the demarcation line is directly related to the concentration of phage. Similar results were found with lysogenic phage lambda and ϕ80. Furthermore, we found commensal *E. coli* K-12 recognizes the pathogen EHEC as non-kin; this recognition has not been identified previously as it was reported that *E. coli* K-12 is not sensitive to the phage of EHEC^18^. Hence, cells employ both active (lytic and temperate) phages and inactive phages (cryptic prophage CPS-53) to control cell to cell contact of commensal gastrointestinal tract bacteria. Although it is generally understood that phage lyse related bacteria that lack the phage, we argue here, that the new phage SW1 is not mere contaminant but instead the *E. coli* K-12 strain uses the phage to its benefit in a social sense (i.e., during nutrient foraging).

**Fig. 1.**
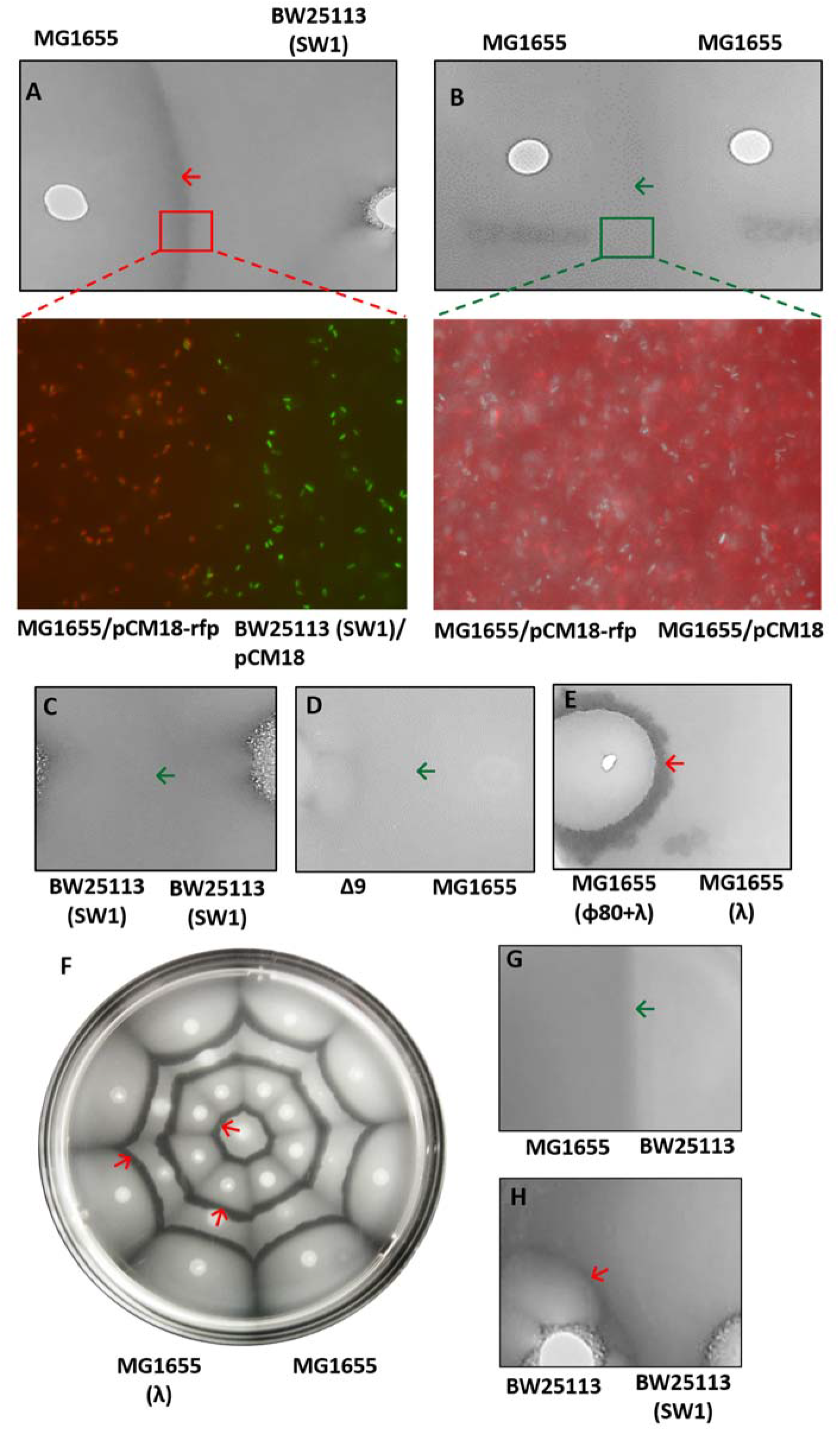
Demarcation lines develop between non-kin for swimming cells. **(A)** Upper panel: Motility halos or MG1655 vs. BW25113 (SW1), lower panel shows non-mixing for non-kin (MG1655/pCM18-rfp appearing as red cells vs. BW25113 (SW1)/pCM18 appearing as green cells). **(B)** Upper panel: MG1655 vs. MG1655, lower panel show mixing for kin (MG1655/pCM18-rfp appearing as red cells vs. MG1655/pCM18 appearing as green cells). Merging of two swimming motility halos for kin **(C)** BW25113 (SW1) vs. BW25113 (SW1), **(D)** BW25113 (SW1) converted to kin by deleting all the cryptic prophage (BW15113 (SW1) Δ9, “Δ9”) vs. MG1655. Demarcation line from the intersection of two swimming motility halos for non-kin **(E)** MG1655 with phages ϕ80 + lambda vs. MG1655 with phage lambda, and **(F)** alternating inoculations of MG1655 with phage lambda (center) and without phage lambda (next concentric circle with eight inoculations). Motility halos for **(G)** MG1655 vs BW25113, and **(H)** BW25113 vs BW25113 (SW1). Each plate contains two strains, and the green arrow indicates where the two swimming halos merge while the red arrow indicates the demarcation line.

## Results

### Non-kin bacteria form demarcation lines during swimming motility

We initially noticed a clear demarcation line between two different *E. coli* K-12 strains, MG1655 and BW25113 (SW1), when inoculated on the same motility plate (**Fig. 1A**); the demarcation line was absent for identical clones: BW25113 (SW1) vs. BW25113 (SW1) and MG1655 vs. MG1655 (**Fig. 1BC**). Hence, the demarcation line was only developed between non-identical strains.

To confirm the motility of identical cells was unaffected when their swimming halos meet, we labeled kin (e.g., MG1655 vs. MG1655) with either RFP and GFP and found complete mixing of cells in the demarcation line (**Fig. 1A**). In contrast, for non-identical strains (i.e., MG1655 vs. BW25113 (SW1)), we found that when their motility halos meet, there was no mixing of cells and that a clear gap formed (**Fig. 1B**).

### The demarcation line between kin is related to phage

We hypothesized the demarcation line was due to a secreted cell product; hence, we searched the complete *E. coli* K-12 library of single-gene knockouts (Keio collection) as a pooled library by plating approximately 10 colonies per motility plate, challenging with MG1655, and looking for the absence of the demarcation line as in indicator of a missing secreted protein. This screen identified 11 mutants (*yahO, nikR, yghG, rpoC, yecA, yfdC, ygeA, mhpA*, and *rep*) that had diminished demarcation lines (**Supplemental Fig. 1**). Of these, only the *rep* mutation completely eliminated the demarcation line. The *rep* mutation has been linked to a swimming and swarming defect previously^19^, and Rep is required for replication of some bacteriophages as an ATP-dependent helicase^20^. Therefore, we focused on phage-related proteins as necessary for the cell to make the demarcation line.

To efficiently check for relationship between the resident *E. coli* cryptic phage and the demarcation line, we tested the Δ9 strain that lacks all nine of the cryptic prophage^12^ vs. MG1655 and found the two motility halos merged and the demarcation line was eliminated (**Fig. 1D**); hence, phage-related proteins are responsible for the demarcation line and self-discrimination. We then tested each of the nine strains with a single, complete cryptic prophage deletion (ΔCP4-6, ΔDLP12, Δe14, Δrac, ΔQin, ΔCP4-44, ΔCPS-53, ΔCPZ-55, and ΔCP4-57)^12^ and found that strains lacking CPS-53 and CP4-57 failed to make a clearance zone (indicating phage lysis) beyond their colonies on a motility plate (**Fig. 2A**) and that deleting CPS-53 prevented BW25113 (SW1) from forming a clearance zone at the boundary of its colony (**Fig. 2A**), whereas, some clearance at the colony boundary was seen for the strain that lacked CP4-57 (**Fig. 2A**). Hence, proteins related to cryptic prophage CPS-53 are required for the demarcation line.

**Fig. 2.**
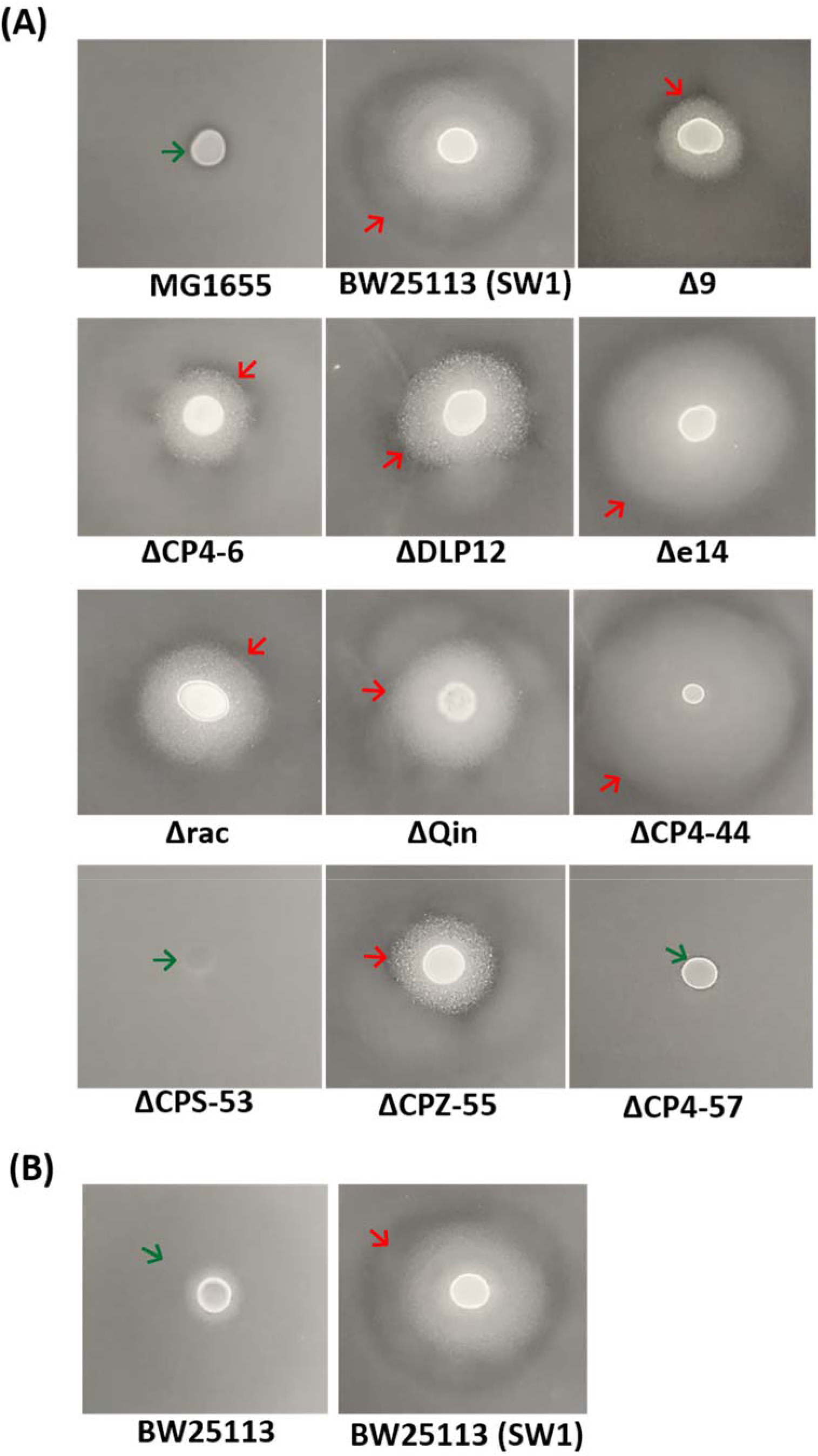
BW25113 cryptic prophage CPS-53 is required to form active phage. Swimming motility halos **(A)** for the strain lacking all nine cryptic prophage, BW15113 (SW1) Δ9 (“Δ9”) and each of the nine BW25113 strains with a single, complete cryptic prophage deletion (ΔCP4-6, ΔDLP12, Δe14, Δrac, ΔQin, ΔCP4-44, ΔCPS-53, ΔCPZ-55, and ΔCP4-57), and **(B)** for BW25113 without SW1 phage and BW25113 (SW1). The green arrow indicates the lack of cell lysis (evidence of phage) while the red arrow indicates the cell lysis. Representative images are shown.

To test whether the complete CPS-53 phage or one its proteins is required to make the demarcation line, we tested 13 single gene mutants of CPS-53 and found that strains with a deletion in CPS-53 prophage genes *yfdQ* and *yfdM* eliminated the demarcation line with MG1655 (**Supplemental Fig. 2**). We then tested these mutants further by trying to restore the demarcation line in these CPS-53 mutants via production of each protein via pCA24N plasmids in its respective BW25113 (SW1) knockout strain and found production of YfdM in the *yfdM* host and YfdQ in the *yfdQ* host re-established the demarcation line with MG1655 (**Supplemental Fig. 2**). Critically, production of these two CPS-53 proteins generated hundreds of plaques with BW25113 (SW1) in the soft agar (**Fig. 3A** for YfdM). We also tested BW25113 (SW1) and found it did not contain lambda phage nor did it contain phage ϕ80 (**Supplemental Fig. 3AB**). In addition, production of YfdM and YfdQ in the ΔCPS-53 strain did not produce plaques, and production of YdfM in the absence of CPS-53 (i.e., ∆CPS-53/pCA24N-yfdM) did not cause a demarcation line to form with MG1655 (**Supplemental Fig. 4A**). These results indicate that cryptic CPS-53 was controlling the production of active SW1 phage particles and that lysis by phage was responsible for producing the demarcation line. This defective-helper relationship has seen be seen previously in EHEC temperate phages^21^.

**Fig. 3.**
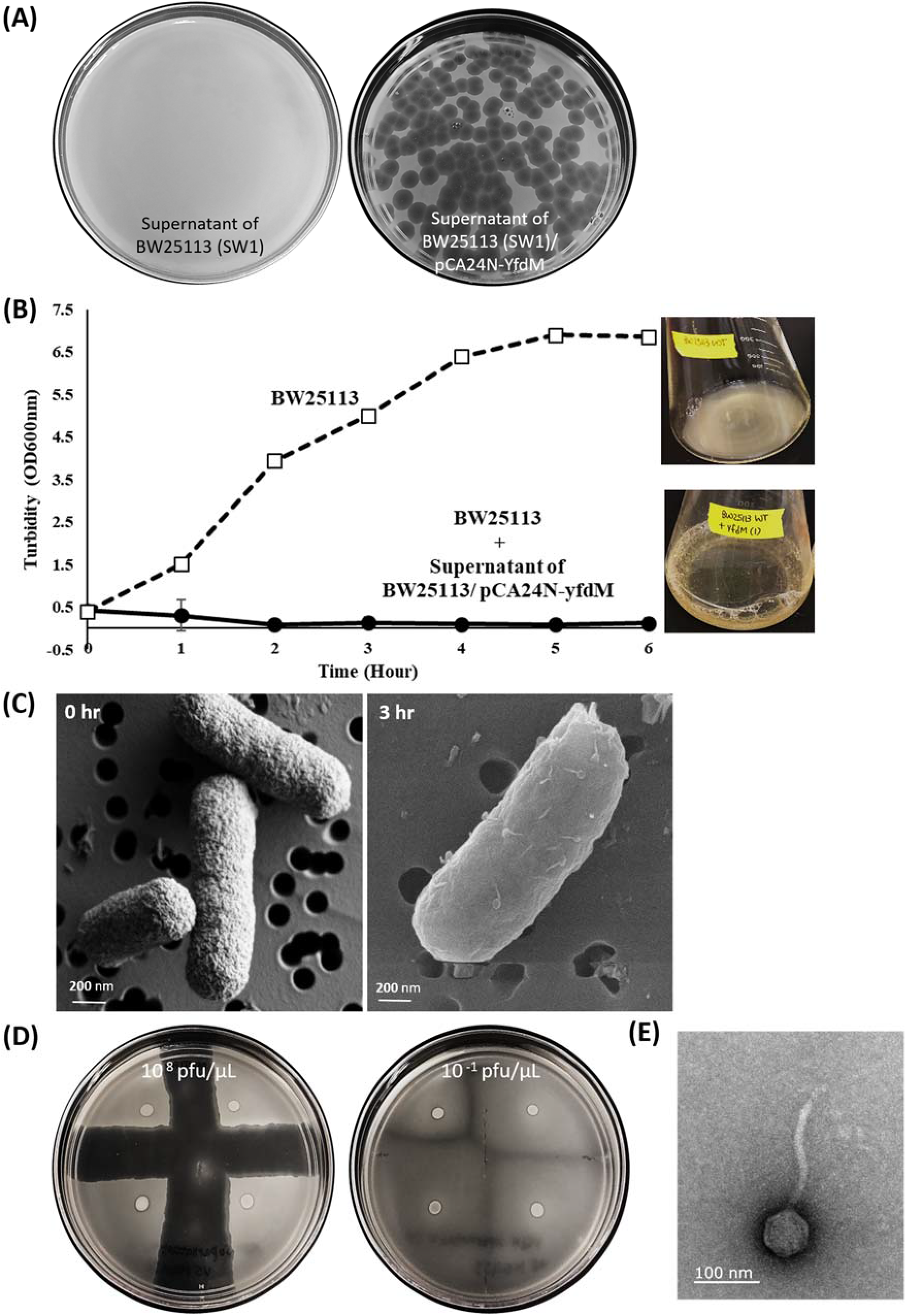
Production of CPS-53 protein YfdM creates active phage. **(A)** Plaque assay ^36^ with BW15113 (SW1) cells in soft agar for the supernatant of BW15113 (SW1) ^36^ and the supernatant of BW25113 (SW1)/pCA24N-yfdM (right) showing plaques are only produced when YfdM is overproduced. **(B)** Production of YfdM in BW25113 (SW1)/pCA24N-yfdM forms supernatants that lyse BW25113 (1% supernatant was added at time 0 and pictures of shake flasks shown as insets with cell debris seen clearly for lysed cells). **(C)** SEM image of the phage particles formed by producing YfdM in BW25113 (SW1)/pCA24N-yfdM that adhere and lyse BW25113 (SW1). Cells shown prior to phage addition (0 h) and after phage build-up (3 h). **(D)** Artificial demarcation line between four MG1655 motility halos created by adding the phage from the supernatant of BW25113 (SW1)/pCA24N-yfdM (left: 10^8^ pfu/uL; right: 10^−1^ pfu/uL). **(E)** TEM image of the phage particles formed by producing YfdM in BW25113/pCA24N-yfdM (amplified 20,000 fold). One representative image is shown.

### Lytic phage SW1 creates the demarcation line

Since CPS-53 is a cryptic prophage unable to produce phage particles, the cell lysis causing the demarcation line must be from some other active phage. To identify the active phage, we produced YfdM from pCA24N-yfdM in BW25113 (SW1) which caused the cell to produce active, lytic phage; i.e., supernatants of BW25113 (SW1)/pCA24N-yfdM induced almost complete cell lysis in the BW25113 (SW1) cells as seen by a dramatic reduction in turbidity of cultures (**Fig. 3B**). We then used scanning electron microscopy to obtain images of the cells at 0 h and 3 h and found the turbidity reduction was caused by phage particles that are seen clearly adhered BW25113 (SW1) cells (**Fig. 3C**). To investigate whether the amount of phage was related to the thickness of the demarcation line, we made an artificial demarcation line with various concentrations of supernatant of BW25113 (SW1)/pCA24N-yfdM containing phage. The supernatant containing phage (10^8^ pfu/μL) formed a thicker barrier in a streak test than supernatants with less phage (10^−1^ pfu/μL) (**Fig. 3D)**, confirming that it was phage causing the demarcation line. Further confirmation of phage causing the demarcation line was that we were able to capture transmission electron micrograph images of the phage made from the supernatants of BW25113 (SW1)/pCA24N-yfdM (**Fig. 3E**); these images indicate production of YfdM causes the production of phage. The phage (SW1) formed round plaques 5 mm in diameter on a lawn of BW25113 (SW1) cells (**Fig. 3A**). The phage has a capsid with a diameter of approximately 71 nm and a non-contractile tail of approximately 205 nm; hence, it is a family siphoviridae phage based on its morphology (**Fig. 3E**).

By sequencing the DNA of the phage particles, we determined SW1 (49,096 bp, **Supplemental Table 3, Supplemental Fig. 5**) is similar to lytic phage T1 (85% identity, 48,836 bp, GenBank AY216660) and phage vB_EcoS_SH2 (99% identity, 49,088 bp, GenBank Y985004). PCR of the three SW1 phage genes (exonuclease, tail fiber, and recombinase) confirmed that SW1 is not present in the BW25113 (SW1) or MG1655 genome; hence, it is a lytic phage (so it does not exist in the genome as a lysogen but exists extracellularly) (**Supplemental Fig. 3C**). Corroborating this, both an integrase and excisionase usually associated with lysogeny are not present in the SW1 genome (**Supplemental Table 3**). Further evidence that SW1 is similar to T1 was found by selecting strains from a pooled Keio collection that could grow in the presence of SW1 phage particles. We found that BW25113 (SW1) cells with a *fhuA* or *tonB* mutation were immune to SW1; FhuA is the attachment protein for T1^22, 23^, and TonB is necessary for transfer through the membrane^22, 23^. Hence, SW1 is a T1-type, lytic phage.

### Lysogenic phages lambda and ϕ80 create a demarcation line

Although our BW25113 (SW1) and MG1655 strains in the preceding results lacked phage lambda and ϕ80, to corroborate the formation of the demarcation line by lysis by SW1, we tested a MG1655 strain that harbored phage lambda. When MG1655 lacking lambda was placed on motility plates with MG1655 containing lambda, a clear demarcation line was formed (**Fig. 1F**). In addition, MG1655 with lysogenic lambda phage formed a double demarcation line with MG1655 with lysogenic ϕ80 (**Fig. 1E**). Hence, demarcation lines are formed from cell lysis due to phage when one of the strains lacks the phage, and two phages cause a more extensive demarcation line to form. To corroborate this result, we tested whether BW25113 lacking SW1 phage (obtained from Coli Genetic Stock Center) forms a demarcation line with MG1655 and found that without SW1, BW25113 does not form a demarcation line with MG1655 **(Fig. 1G)**. As expected, BW25113 forms a demarcation line with BW25113 harboring the SW1 phage **(Fig. 1H).**

### Colonies with phage form a colony demarcation line on motility plates

Observation of colonies on motility plates provided additional evidence of the presence of phage and cell lysis; for example, BW25113 lacking SW1 does not show evidence of cell lysis around its colony, but BW25113 infected with SW1 shows a demarcation line that forms beyond the colony and surrounds it (**Fig. 2B**); we termed this demarcation line around a colony as a “colony demarcation line”. We found this colony demarcation line phenomenon for MG1655 including phages lambda and ϕ80 while the MG1655 lacking phage does not show this phenotype (**Supplemental Fig. 6A**). In addition, MG1655 infected by SW1 phage also has a colony demarcation line (**Supplemental Fig. 6A**). These results demonstrate that the presence of active phage can be discerned from the colony demarcation line for colonies on motility plates.

Moreover, using the colony demarcation line approach, we found that colonies of BW25113 ∆*yfdM* and BW25113 ∆CPS-53 were not changed by SW1 phage infection (**Supplemental Fig. 7**). Hence, the absence of the colony demarcation line shows again the dependence of SW1 on proteins of cryptic prophage CPS-53, specifically, the dependence on YfdM.

### Cells lyse in the demarcation line

To determine the role of phage in kin recognition, we investigated whether cells lyse in the demarcation line when phage are produced. We found that although the demarcation line appears clear on motility plates, there are some cells in this zone, and their numbers are about 10-fold less than in the periphery of the swimming halo (**Fig. 4A**, **Supplemental Table 4**). Critically, we found that for strains that produce a demarcation line, i.e., for MG1655 vs. BW25113 (SW1), cells are dead in the demarcation line (**Fig. 4AB**) which indicates phage propagation (i.e., lysis) throughout its swimming halo. In contrast, for strains not producing a demarcation line; i.e., for BW25113 (SW1)/pCA24N vs. BW25113 (SW1)/pCA24N (**Fig. 4C**), all the cells were healthy and unlysed in the merging motility halos. These results demonstrate clearly that the demarcation line is formed from the lysis of non-kin and that kin have some immunity from cell lysis by phage SW1 when swimming cells meet.

**Fig. 4.**
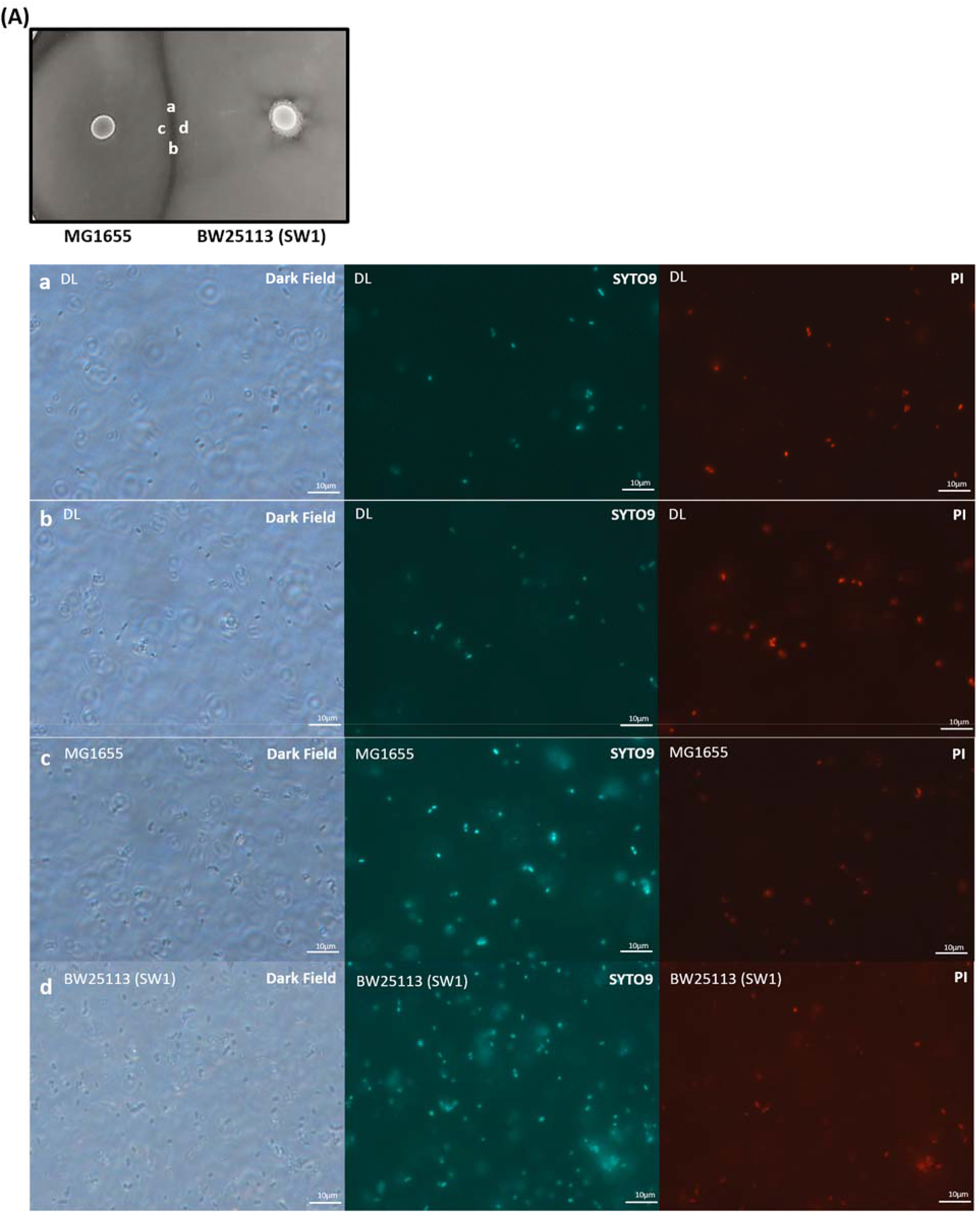
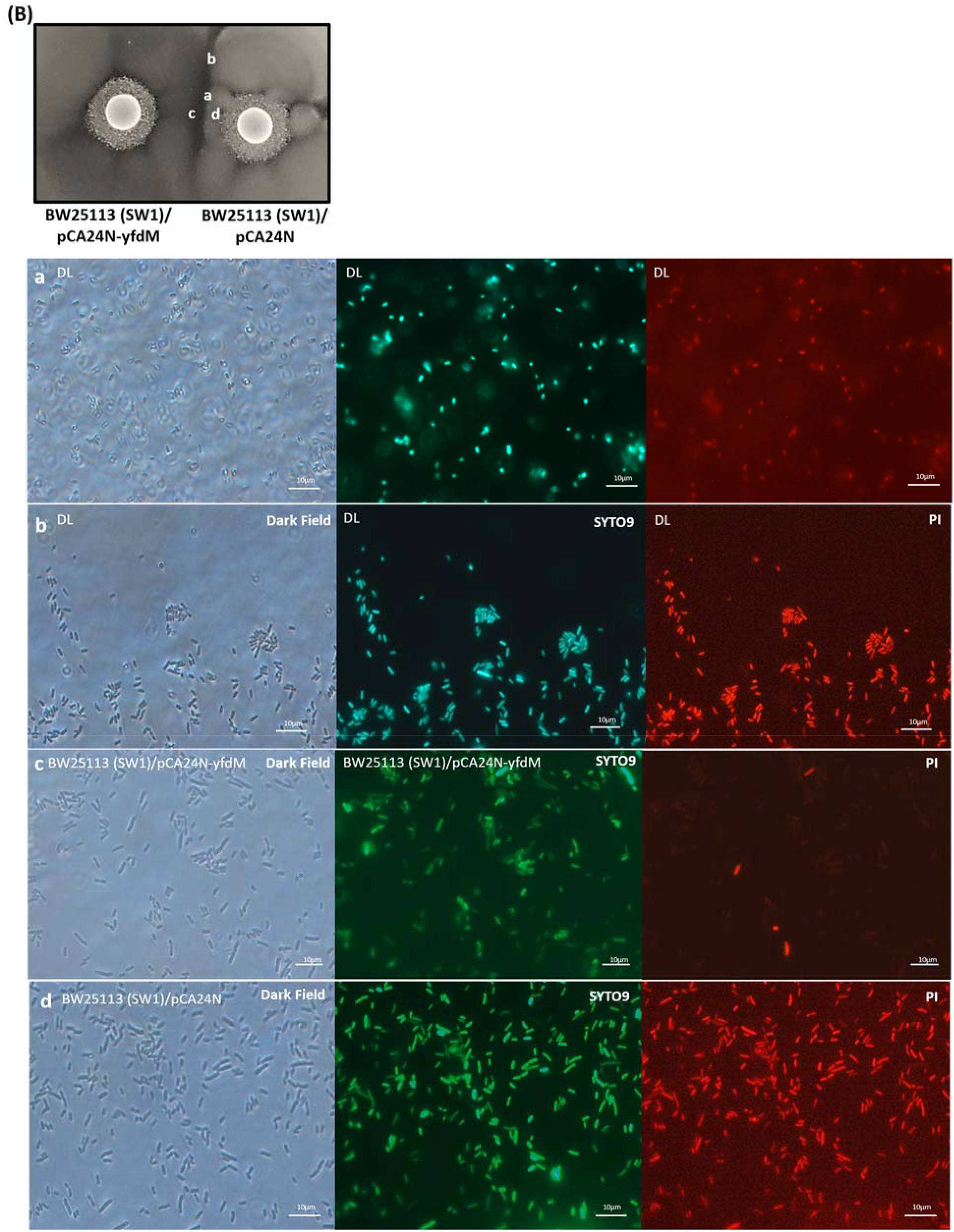
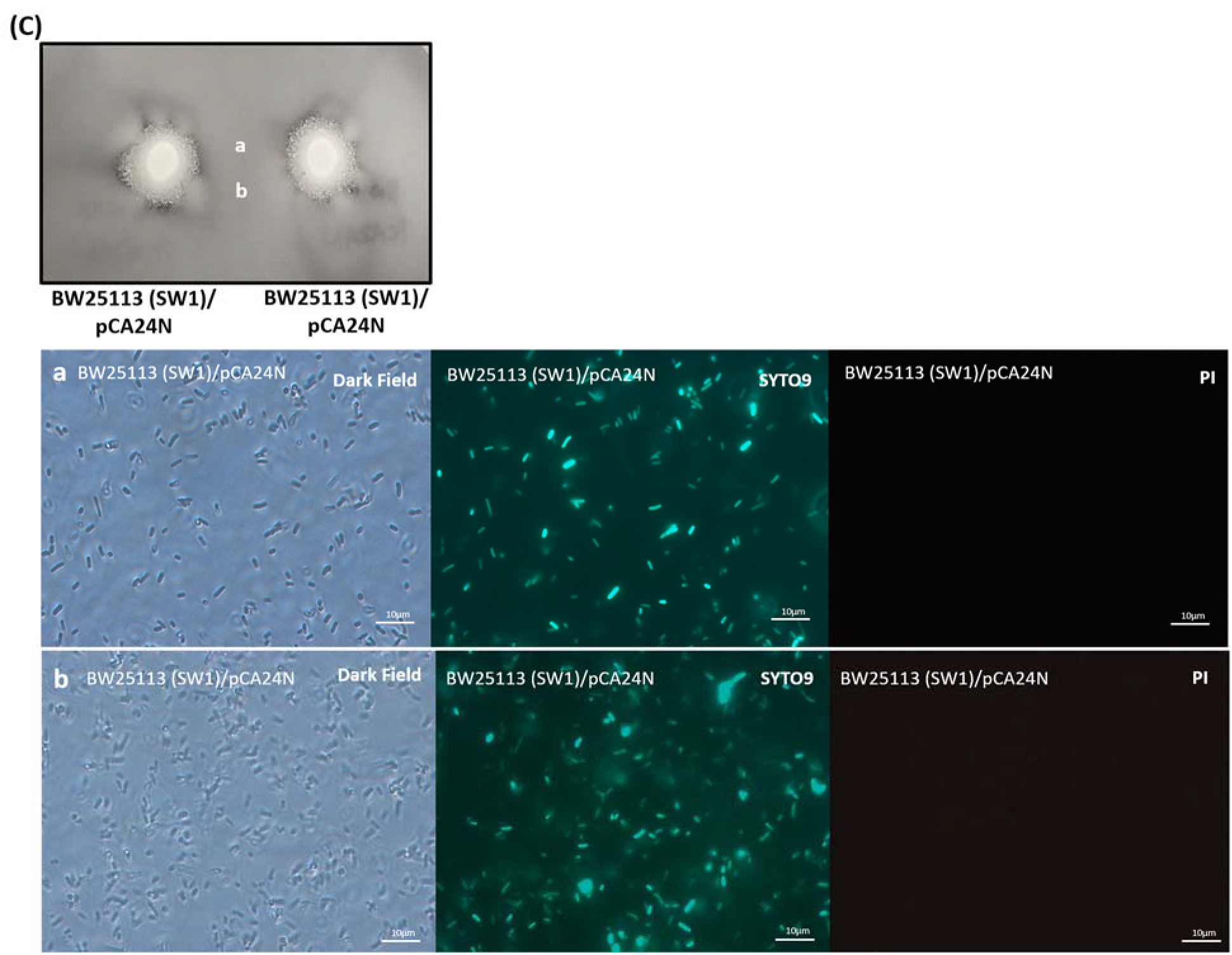
Cell death causes the demarcation line. Live/Dead staining of the intersection of the swimming motility halos of **(A)** MG1655 vs. BW25113 (SW1), **(B)** BW15113 (SW1)/pCA24N-yfdM vs. BW25113/pCA24N, and **(C)** BW15113 (SW1)/pCA24N vs. BW15113 (SW1)/pCA24N. Upper panels: photo letters “a”, “b”. “c”, and “d” indicate the position imaged in the fluorescence microscope. Lower panels: left shows dark field, middle shows all cells stained by Syto 9, and right shows dead cells stained by propidium iodide. Representative images are shown.

### YfdM together with phage SW1 can convert kin to non-kin

Based on these results, we reasoned that cells could be made to no longer recognize kin if one strain over-produced phage. Verifying this hypothesis, overproduction of CPS-53 phage protein YfdM in BW25113 to cause the overproduction of phage caused the formation of a very large demarcation line with BW25113 (SW1)/pCA24N (**Fig. 5A**) in which cells were clearly killed. In contrast, without YfdM production, the two BW25113 (SW1) strains merged perfectly (**Fig. 5B**) and production of YfdM in the absence of CPS-53 from both strains was unable to produce a demarcation line **(Fig. 5C)**. Additionally, production of YfdM in MG1655 (i.e., MG1655/pCA24N-yfdM) was unable to produce a demarcation line with MG1655/pCA24N (**Fig. 5D**), since MG1655 lacks phage SW1. As expected, two MG1655/pCA24N strains merge perfectly (**Fig. 5E)**. Therefore, YfdM together with SW1 are necessary to form the demarcation line in BW25113, and kin can be converted to non-kin by the production of phage.

**Fig. 5.**
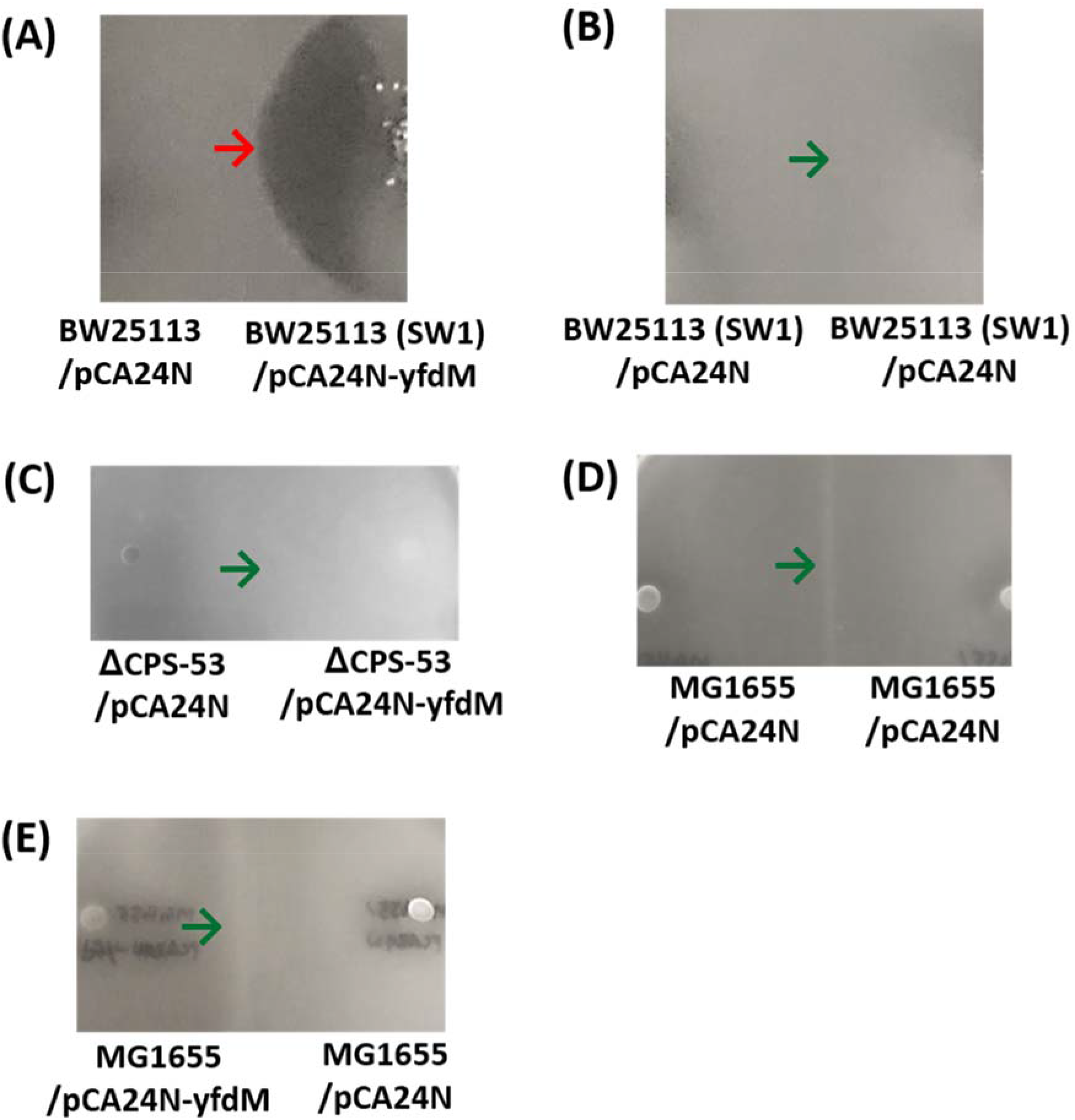
Production of CPS-53 protein YfdM creates a sharp demarcation line in the presence of SW1. Intersection of swimming motility halos of **(A)** BW25113 (SW1)/pCA24N vs. BW25113 (SW1)/pCA24N-yfdM, **(B)** BW15113 (SW1)/pCA24N vs. BW15113 (SW1)/pCA24N, **(C)** MG1655/pCA24N-yfdM vs. MG1655/pCA24N, **(D)** MG1655/pCA24N vs. MG1655/pCA24N, and **(E)** ∆CPS-53/pCA24N vs. ∆CPS-53/pCA24N-yfdM. YfdM was produced from pCA24N-yfdM with 0.5 mM IPTG. Each plate contains two strains, and the green arrow indicates where the two swimming halos merge while the red arrow indicates the demarcation line. Representative images are shown. Note the lines seen for panels C and D are due to the accumulation of cells at the place where the motility halos combine.

To confirm that kin can be converted to non-kin through phage, we infected MG1655 with SW1 phage, and tested whether demarcation line could be created with MG1655 that lack phage SW1. We found MG1655 with phage SW1 forms a large demarcation line with MG1655 that lacks SW1; hence, again, phage can be used to convert kin to non-kin (**Supplemental Fig. 6B**).

### Gastrointestinal recognition through phage

Additionally, we investigated whether commensal K-12 could recognize pathogenic bacteria such as EHEC through the use of phage. We found K-12 formed a demarcation line with EHEC (**Supplemental Fig. 8**); in contrast, the swimming halos of two EHEC colonies merged. These results confirm kin are recognized and non-kin are lysed with pathogenic bacteria. In addition, EHEC colonies have a colony-demarcation line (**Supplemental Fig. 8J**), and EHEC swimming cells form a smaller but discernable demarcation line with the K-12 strains that lack SW1 (**Supplemental Fig. 8C vs. 8D**) indicating EHEC also uses its own active phage particles^21^ for kin recognition. Further proof that EHEC phage recognizes *E. coli* K-12 is that we found that phage particles isolated from EHEC form plaques with both BW25113 (SW1) and BW25113 that lacks SW1 (**Supplemental Fig. 8K, L).** Hence, EHEC competes with *E. coli* K-12.

### SW1 increases cell fitness

To determine whether SW1 increased cell fitness, we labelled cells to distinguish their motility halos and monitored swimming motility for BW25113 (SW1) vs. BW25113. In six out of seven trials, cells associated with SW1 were able to engulf those that lack SW1 showing the lytic phage SW1 provides a demonstrable benefit during nutrient foraging (**Fig. 6A**). Independently, the motility of the two strains was not significantly different (**Fig. 6BC**).

**Fig. 6.**
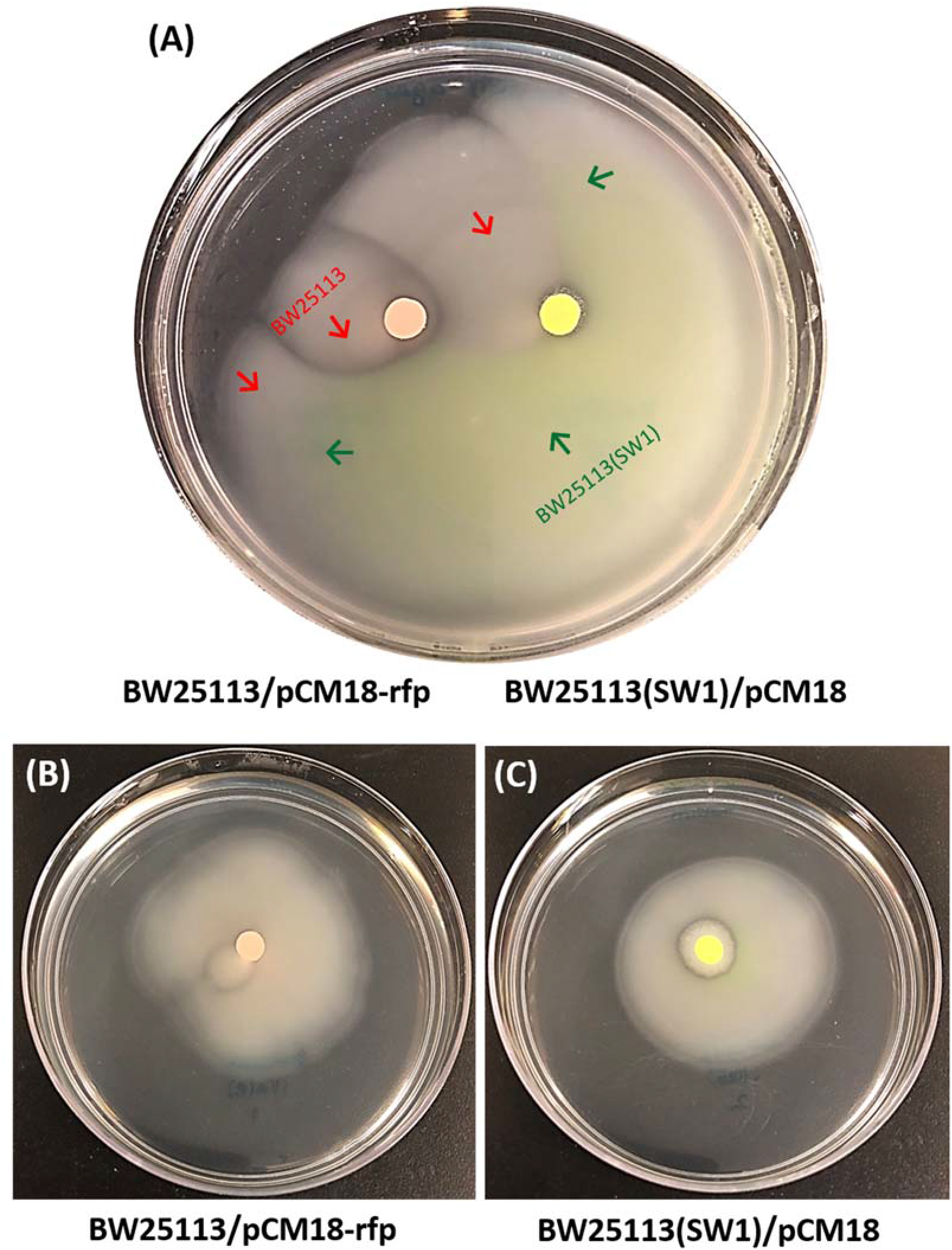
SW1 increases the competitiveness of BW25113 during nutrient foraging. **(A)** Swimming motility of BW25113/pCM18-rfp (light pink from RFP) vs. BW25113 (SW1)/pCM18 (yellow-green from GFP) after 28 h. The plate contains two strains, and the green arrows indicate the halo of BW25113 (SW1)/pCM18, which contains lytic phage SW1, while the red arrows indicate the halo of BW25113/pCM18-rfp, which lacks SW1. One representative image from seven independent cultures shown. Individual swimming motility of **(B)** BW15113/pCM18-rfp and **(C)** BW25113(SW1)/pCM18 after 28 h was not significantly different. One of seven independent cultures shown.

### Origin of phage SW1

To determine the source of the SW1 phage, we used the colony demarcation line assay for nine random Keio mutants since we originally found the colony demarcation line in our BW25113 wild-type stock from the Keio collection^17^. We found colony demarcation lines in five Keio mutants from the original source plates (**Supplemental Fig. 9**), and three of these strains, 15-G9, 15-E6, and 63-D7 had phage (30 pfu/100 μL) whose numbers increased by 10-fold with YfdM production. Also, we checked hundreds of Keio mutants for the Dienes line assay with MG1655, and most of the mutants formed a demarcation line indicating the presence of SW1 (**Supplemental Fig. 1**). Hence, phage SW1 may be pervasive through dissemination of the Keio collection (and visible in specific clones via the colony demarcation assay). Note full activation of SW1 (i.e., full cell lysis) requires electroporation of putative Dam methylase YfdM of cryptic prophage CPS-53 (e.g., pCA24N-yfdM).

## Discussion

We demonstrate here for the first time that cells use phage to discriminate between kin and non-kin. Hence, during motility, when two strains meet and one strain lacks a phage to which it is sensitive, a clearance zone forms in which the cells that lack the phage are lysed (**Fig. 4**). Although we did not investigate the role of phage in biofilms (we focused on swimming motility), we anticipate that similar principles apply and that phage-containing cells have a fitness advantage over non-phage containing cells.

We also demonstrate here that commensal *E. coli* uses cell killing based on proteins from cryptic prophage CPS-53 (i.e., YfdM) to activate lytic phage SW1 to differentiate itself from non-kin. Hence, our results show a surprising dependence of lytic phage SW1 on cryptic prophage proteins; this result provides a new insight into the benefit of cells harboring cryptic prophage. Of course, dependence on chromosomal proteins (i.e., non-cryptic prophage proteins) has been demonstrated previously with T1-type phage; for example, as we found here, FhuA is required for attachment and TonB is required for transfer phage through the cell membrane^22, 23^. Critically, YfdM of CPS-53 has 91 aa in common with the Dam methylase of EHEC Stx2 phage^24^, and T1 requires Dam. Moreover, putative methylase YfdM is not related to the Dam methylase of SW1 or BW25113, so it should provide a novel methylase function. Hence, SW1 appears to use the putative Dam methylase of CPS-53 (i.e., YfdM) to make active phage. Supporting this hypothesis, bacteriophage P1 cleaves its DNA at a packing site (*pac*) once it is methylated by adenine methylase^25, 26^, and T1 also has a *pac* site^27^. Hence, YfdM of CPS-53 might methylate a *pac-*like site of SW1 to signal DNA cleavage for filling the prohead with DNA during phage packaging, and this methylase activity may limit DNA packaging^27^. Evidence of this is that we only see perfectly round plaques in the presence of overproduced YfdM (**Fig. 3A**). Alternatively, putative methylase YfdM may facilitate SW1 proliferation by protecting it from restriction enzyme attack. Notably, we only obtain a large number of SW1 phage particles when YfdM is overproduced, but SW1 propagates whenever there is a wild-type copy of *yfdM^+^* in the chromosome.

Since some cell death occurs in the absence of neighboring cells (see the lysis zones we named colony demarcation lines around colonies on motility plates, **Fig. 2**), the use of lytic phage requires some cell altruism as there is clearly some cell death for the host using this phage approach. Overall, the use of lytic phage particles should increase the fitness of commensal *E. coli* as long as there is more lysis of the competing cells. What remains to be determined is how kin are protected from phage SW1 since clearly kin have no demarcation line (**Fig. 1BC**); perhaps, its lysis is growth-phase dependent or there is a SW1 protein that delays lysis and gives immunity. Clearly lysogenic immunity is not involved for SW1 as there is no lysogenic phase for this phage. Since for lysogenic phage such as lambda and ϕ80 there should be phage immunity for the phage-bearing strain^28, 29^, the phage-containing strain should have an even greater fitness advantage over the phage-minus strain.

In the gastrointestinal tract, commensal *E. coli* cells must compete with one thousand other bacterial species, whose combined density reaches 10^12^ organisms per gram^6^. In addition, about five different *E. coli* strains usually colonize each human gastrointestinal tract^30^. Furthermore, the chief competitor of pathogenic *E. coli* (EHEC) in the gastrointestinal tract is *E. coli* K-12^18^, and we demonstrated a demarcation line is formed between these two strains based the phages of both EHEC and BW25113 (SW1) since EHEC forms a demarcation line BW25113 that lacks phage (**Supplemental Fig. 8I vs. Supplemental Fig. 8D**). We also found phages from EHEC form plaques on BW25113 (**Supplemental Fig. 8K, L**) and found that the presence of SW1 makes BW25113 more competitive during nutrient foraging (**Fig. 6**). Hence, self-recognition should be paramount to the success of this commensal bacterium in increasing its fitness in colonizing humans as a collective group, since it is a non-dominant strain^31^ and must compete with similar *E. coli* strains.

Note that there are many phages in the gastrointestinal tract^32^, so the use of phage for kin selection is reasonable. This approach for kin recognition is also general based on our results with three phages including both lytic (e.g., SW1) and lysogenic phage (e.g., lambda, ϕ80), and based on our results with both commensal *E. coli* and pathogenic *E. coli.*

## Methods

### Bacterial species, growth conditions, and plasmids

The species and plasmids used in this study are listed in **Supplemental Table 1**. For deleting and overexpressing genes, we used the Keio collection and the ASKA library. All experiments were conducted at 37°C, and cells were routinely cultured in lysogeny broth (LB)^33^. Kanamycin (50 μg/mL) was used for pre-culturing the isogenic knockout mutants, chloramphenicol (30 μg/mL) was used for maintaining the pCA24N-based plasmids, and erythromycin (300 μg/mL) was used for maintaining the pCM18-based plasmids.

### Motility halo assay

Swimming motility was examined using 0.3% agar plates with 1% tryptone and 0.5% NaCl. Motility and demarcation lines were analyzed after 24 h. Three plates were tested for each culture, and two independent cultures were used. Cultures (5 μL) were dropped on motility agar plates and each plate contains two strains for observing demarcation line formation and one strain for observing colony demarcation line formation; plates were incubated 16 h to form motility halos. YfdM was produced via pCA24N-yfdM via its leaky promoter or via 0.5 mM isopropyl β-D-1-thiogalactopyranoside (IPTG) as indicated. The motility agar plates were dried for 2 h to achieve consistent results.

### Screening via the pooled Keio collection

To identify which *E. coli* BW25113 proteins are related to the demarcation line with MG1655, we inoculated the pooled Keio collection onto LB agar plates with kanamycin (50 μg/mL). After incubating at 37°C for 16 h, one colony from the Keio collection was inoculated with MG1655 on the same motility plate; colonies that merged with the MG1655 halo were selected, and the candidates were sequenced with primer KFp1 (**Supplemental Table 2**). The first PCR was performed with primer Arb1 and Inv-2 with 94°C for 5 min, (94°C for 30 sec, 30°C for 30 sec, 72°C for 1 min)*5 cycles, (94°C for 30 sec, 52°C for 30 sec, 72°C for 1 min)*30 cycles, and 72°C for 5 min. The second PCR reaction was performed with the PCR product from the first PCR reaction with primers (Arb2 and Kfp-1) using the conditions: (94°C for 30 sec, 50°C for 30 sec, 72°C for 1 min) *30 cycles.

To identify which proteins are required for SW1 phage infection in BW25113, SW1 phage stock (10^11^ pfu/mL) was added (10%) to the pooled Keio culture with a turbidity of 1.0 at 600 nm. The culture was spread on LB agar plates with kanamycin when the culture reached a turbidity of 0.1 at OD 600, and the DNA from the colonies was sequenced.

### Gel pad microscopy with GFP/RFP-labeled strains

The GFP/RFP labeled strains were inoculated on motility agar on glass slides and incubated overnight. The motility agar was covered with a cover glass, and the fluorescence signal was analyzed by microscopy (Zeiss Axioscope.A1) using excitation at 488 nm and emission at 500 nm for GFP fluorescence and using excitation at 561 nm and emission at 575 nm for RFP fluorescence.

### Plaque assay

Supernatants from the strains were harvested by centrifugation (8000 *g* for 10 min) and filtered (0.2 μm), then phage were quantified by counting plaques using a modified top-layer agar method^34^. The plaque assay was performed with three independent cultures for each strain with three replicates for each culture. EHEC phage were induced by mitomycin C (10 μg/mL) for getting EHEC phage.

### Polymerase chain reaction (PCR) to detect phages

To detect lambda phage and ϕ80, PCR was performed with a colony that was mixed with 20 μL of water and boiled for 10 min; 2 μL of the crude cell extract was used with primers shown in **Supplemental Table 2**. To detect phage genes in SW1, phage DNA was used as the DNA template with primers shown in **Supplemental Table 2** after isolating the phage DNA by the Phage DNA isolation kit 46800 (Norgen Biotetek Corp). To detect SW1 phage genes associated with BW25113 (SW1) and BW25113 (SW1)/pCA24N-yfdM, genomic DNA was used for BW25113 (SW1) and a culture (i.e., cells + supernatant) of BW25113 (SW1)/pCA24N-yfdM was used as the DNA template with primers shown in **Supplemental Table 2.**

### Electron microscopy

For transmission electron microscopy (TEM), the BW25113/pCA24N-YfdM culture was cultured for 48 h to lyse cells, centrifuged at 8000 *g*, and the supernatant was filter-sterilized (0.2 μm). TEM images were obtained using a JEOL JEM 1200 EXII instrument. For scanning electron microscopy (SEM), the samples were collected at 0 h, 3 h, and 6 h after adding the phage stocks to BW25113 cells. The sample was fixed with buffer (2.5% glutaraldehyde in 0.1M cacodylate buffer, pH 7.4). SEM images were obtained using a Zeiss SIGMA VP-FESEM instrument.

### Viability assay

Once the demarcation line between the strains was formed, it was removed from the 300 motility agar and stained for 1 h at room temperature in the dark using the using the LIVE/DEAD BacLight Bacterial Viability Kit L7012 (Molecular Probes, Inc., Eugene, OR). The fluorescence signal was analyzed via microscopy using excitation at 485 nm and emission at 530 nm for green fluorescence and using excitation at 485 nm and emission at 630 nm for red fluorescence (Zeiss Axioscope.A1).

### Infection with SW1 phage

To infect bacteria with SW1 phage, phage stock (2% by vol) was added to a culture when the strain reached a turbidity of 1.0 at 600 nm and it was incubated for 24 h.

### Phage DNA genome sequencing

Phage DNA was isolated from BW25113/pCA24N-YfdM after growth in LB for 36 h by the Phage DNA isolation kit (catalog number 46800, Norgen Biotetek Corp). The DNA phage was sequenced by an Oxford Nanopore Technologies (ONT) MinION^35^ instrument.

## Acknowledgements

We are grateful for the pooled Keio library provided by Dr. Justin Gallivan and for the TEM and SEM images provided by Missy Hazen of the Huck Institutes of the Life Sciences at the Pennsylvania State University. This work was supported by the Army Research Office (W911NF-14-1-0279) and funds derived from the Biotechnology Endowed Professorship at the Pennsylvania State University.

## Author contributions

SYS, YG, and JSK conducted the experiments. XW and TKW conceived the project. SYS and TKW wrote the manuscript.

## Reprints and permission (Data and materials availability)

All data are available in the main text or in the supplementary materials except for the DNA sequence of phage SW1 which has been deposited at NCBI.

## SUPPLEMENTARY MATERIAL

**Supplemental Table 1.** Strains and plasmids used in this study.

| **Strains & Plasmids** | **Genotype/relevant characteristics** | **Source** |
| --- | --- | --- |
| MG1655 | Wild type: F<sup>-</sup> $\lambda^-$ *ilvG*<sup>-</sup> *rfb*-50 *rph*-1 | F. Blattner |
| MG1655 Δ10TA | Δ*mazF* Δ*chpB* Δ*relBE* Δ*dinJ*-*yafQ* Δ*yefM*-*yoeB* Δ*higBA* Δ*prlFyhaV*<br>Δ*yafNO* Δ*mqsRA* Δ*hicAB* λdef(+) ϕ80(+) ϕ80h(80)imm($\lambda$)(+) | <sup>37</sup> |
| MG1655 $\lambda$+ | Wild type including lambda phage *lamB* $\lambda$(+) | lab stock |
| O157:H7 EDL933 | Wild type EHEC | <sup>38</sup> |
| BW25113 | Wild type: *lacI*<sup>q</sup> *rrnB*<sub>T14</sub> Δ*lacZ*<sub>WJ16</sub> *hsdR*514 Δ*araBAD*<sub>AH33</sub> Δ*rhaBAD*<sub>LD78</sub> | Coli Genetic Stock Center |
| BW25113 (SW1) | Wild type including SW1 phage | <sup>17</sup> |
| **BW25113 derivatives** |  |  |
| *yahO* | Δ*yahO* ΩKm<sup>R</sup> | <sup>17</sup> |
| *nikR* | Δ*nikR* ΩKm<sup>R</sup> | <sup>17</sup> |
| *yghG* | Δ*yghG* ΩKm<sup>R</sup> | <sup>17</sup> |
| *rpoC* | Δ*rpoC* ΩKm<sup>R</sup> | <sup>17</sup> |
| *insQ* | Δ*insQ* ΩKm<sup>R</sup> | <sup>17</sup> |
| *yecA* | Δ*yecA* ΩKm<sup>R</sup> | <sup>17</sup> |
| *appY* | Δ*appY* ΩKm<sup>R</sup> | <sup>17</sup> |
| *yfdC* | Δ*yfdC* ΩKm<sup>R</sup> | <sup>17</sup> |
| *ygeA* | Δ*ygeA* ΩKm<sup>R</sup> | <sup>17</sup> |
| *mhpA* | Δ*mhpA* ΩKm<sup>R</sup> | <sup>17</sup> |
| *rep* | Δ*rep* ΩKm<sup>R</sup> | <sup>17</sup> |
| *yfdR* | Δ*yfdR* ΩKm<sup>R</sup> | <sup>17</sup> |
| *yfdG* | Δ*yfdG* ΩKm<sup>R</sup> | <sup>17</sup> |
| *yfdL* | Δ*yfdL* ΩKm<sup>R</sup> | <sup>17</sup> |
| *intS* | Δ*intS* ΩKm<sup>R</sup> | <sup>17</sup> |
| *yfdQ* | Δ*yfdQ* ΩKm<sup>R</sup> | <sup>17</sup> |
| *yfdS* | Δ*yfdS* ΩKm<sup>R</sup> | <sup>17</sup> |
| *yfdM* | Δ*yfdM* ΩKm<sup>R</sup> | <sup>17</sup> |
| *yfdI* | Δ*yfdI* ΩKm<sup>R</sup> | <sup>17</sup> |
| *yfdR* | Δ*yfdM* ΩKm<sup>R</sup> | <sup>17</sup> |
| ΔCP4-57 | BW25113 ΔCP4-57 | <sup>12</sup> |
| Δrac | BW25113 Δrac | <sup>12</sup> |
| Δe14 | BW25113 Δe14 | <sup>12</sup> |
| ΔCPS-53 | BW25113 ΔCPS-53 | <sup>12</sup> |
| ΔCP4-6 | BW25113 ΔCP4-6 | <sup>12</sup> |
| ΔDLP12 | BW25113 ΔDLP12 | <sup>12</sup> |
| ΔQin | BW25113 ΔQin | <sup>12</sup> |
| ΔCP4-44 | BW25113 ΔCP4-44 | <sup>12</sup> |
| ΔCPZ-55 | BW25113 ΔCPZ-55 | <sup>12</sup> |
| Δ9 | ΔCP4-57 Δrac Δe14 ΔCPS-53 ΔCP4-6 ΔDLP12 ΔQin ΔCP4-44 ΔCPZ55 | <sup>12</sup> |
| **Plasmids** |  |  |
| pCA24N | Cm<sup>R</sup>; *lacI*<sup>q</sup> | <sup>39</sup> |
| pCA24N-yfdM | Cm<sup>R</sup>; *lacI*<sup>q</sup>, P<sub>T5-lac</sub>::*yfdM*<sup>+</sup> | <sup>39</sup> |
| pCA24N-yfdQ | Cm<sup>R</sup>; *lacI*<sup>q</sup>, P<sub>T5-lac</sub>::*yfdQ*<sup>+</sup> | <sup>39</sup> |
| pCM18 | Em<sup>R</sup>; P<sub>CP25</sub>::*gfp*<sup>+</sup> | <sup>40</sup> |
| pCM18-rfp | Em<sup>R</sup>; P<sub>CP25</sub>::*gfp*<sup>+</sup>-*rfp*<sup>+</sup> | <sup>41</sup> |

**Supplemental Table 2.**
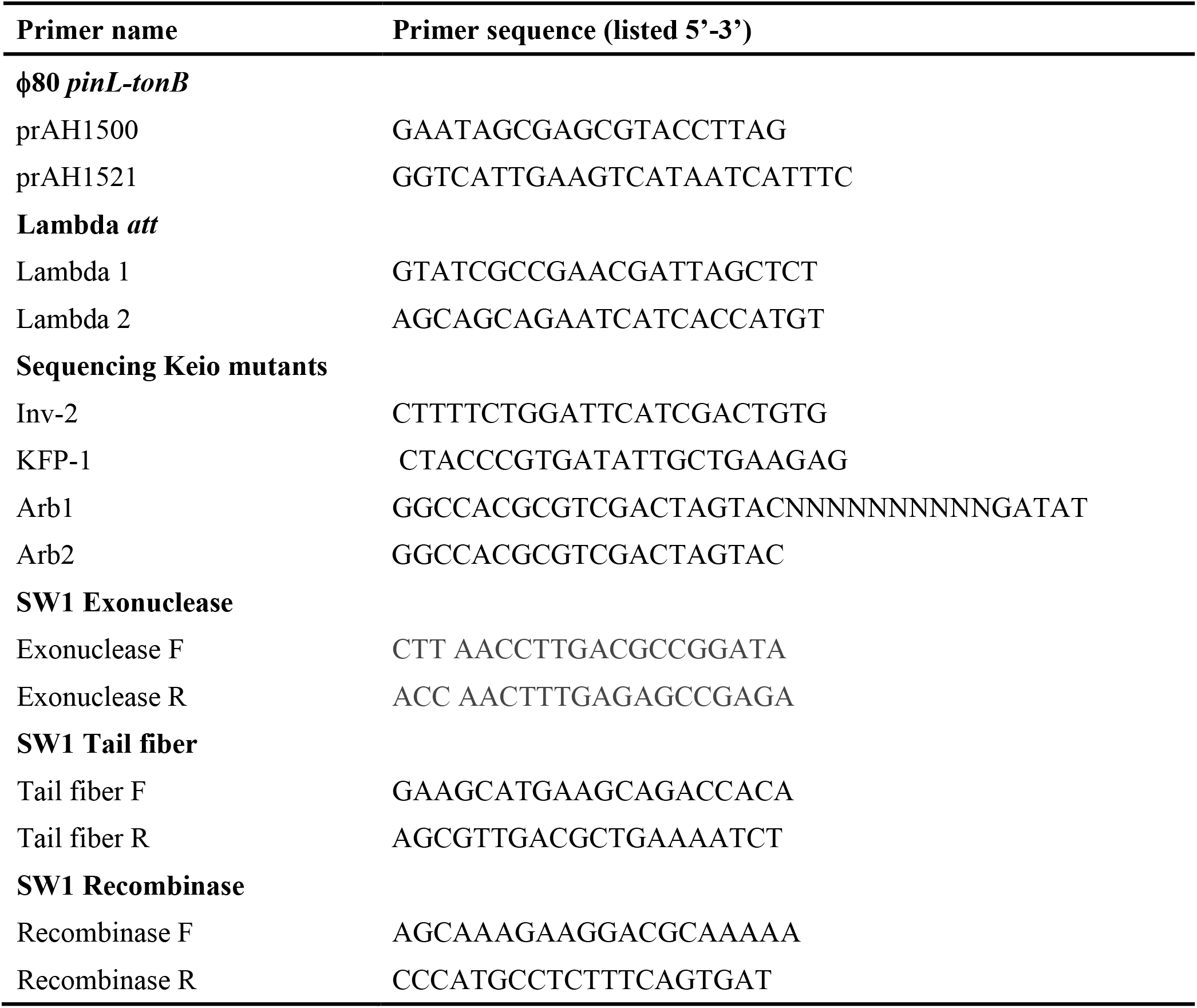
Primers used for PCR.

**Supplemental Table 3.**
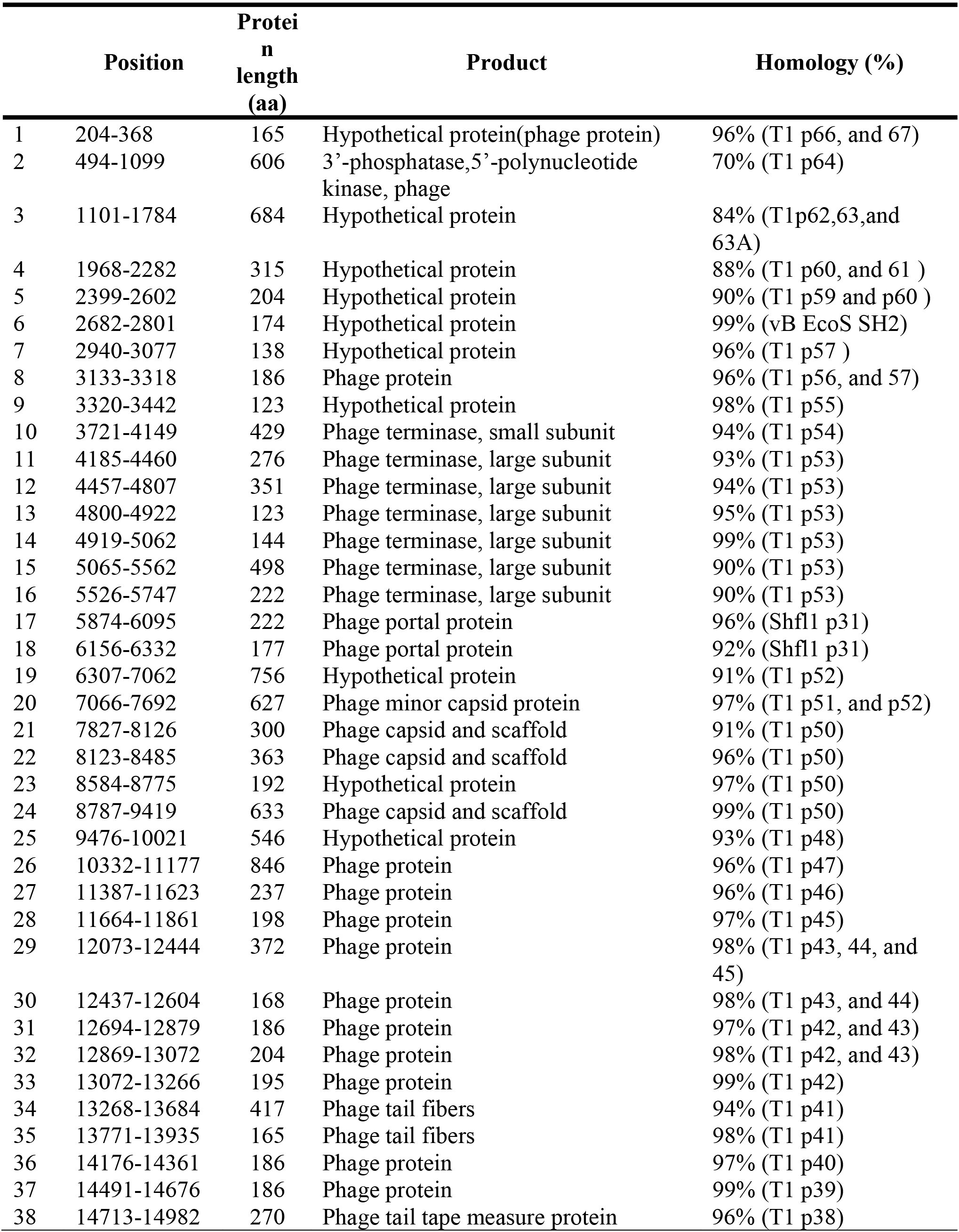
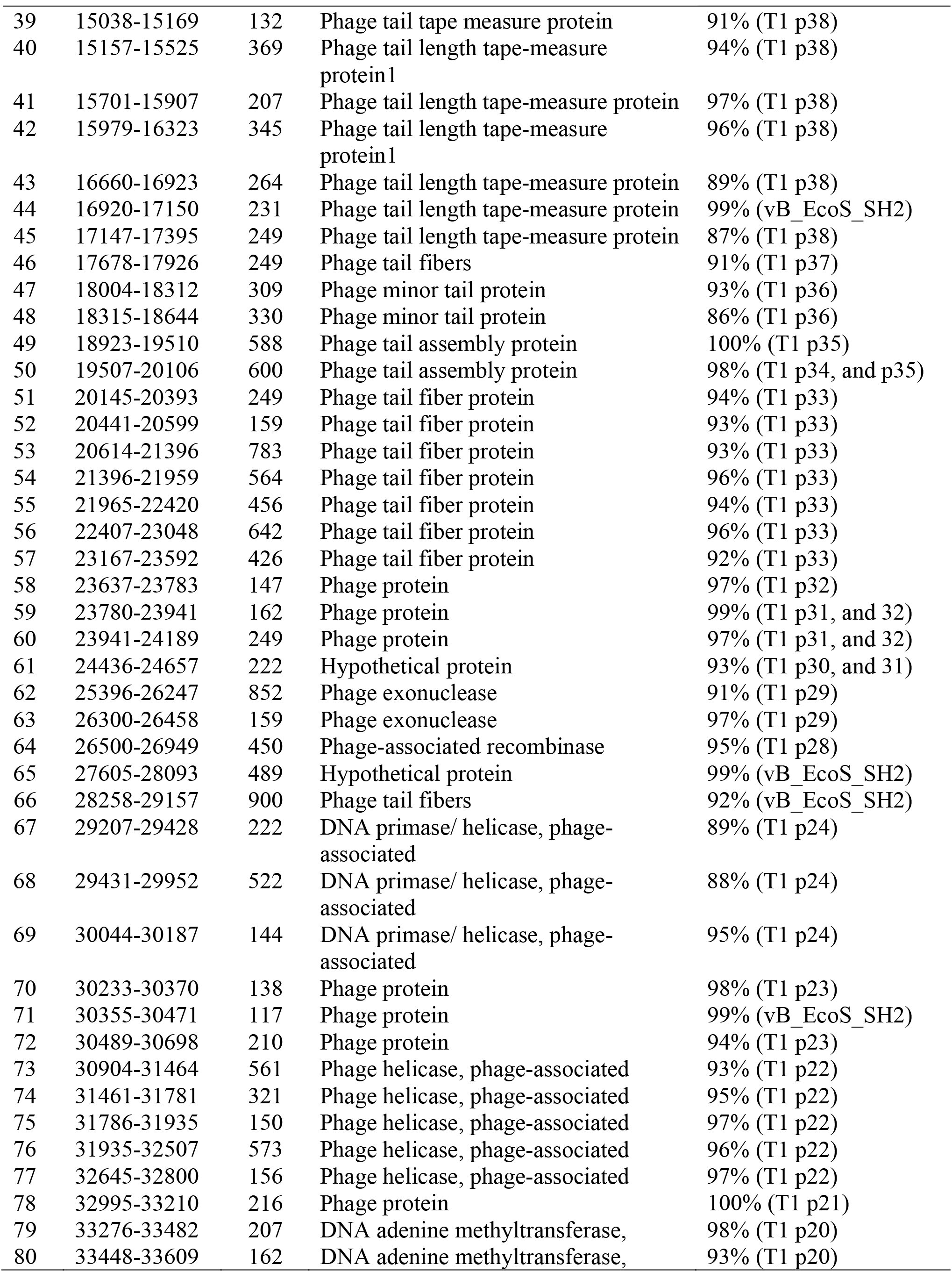
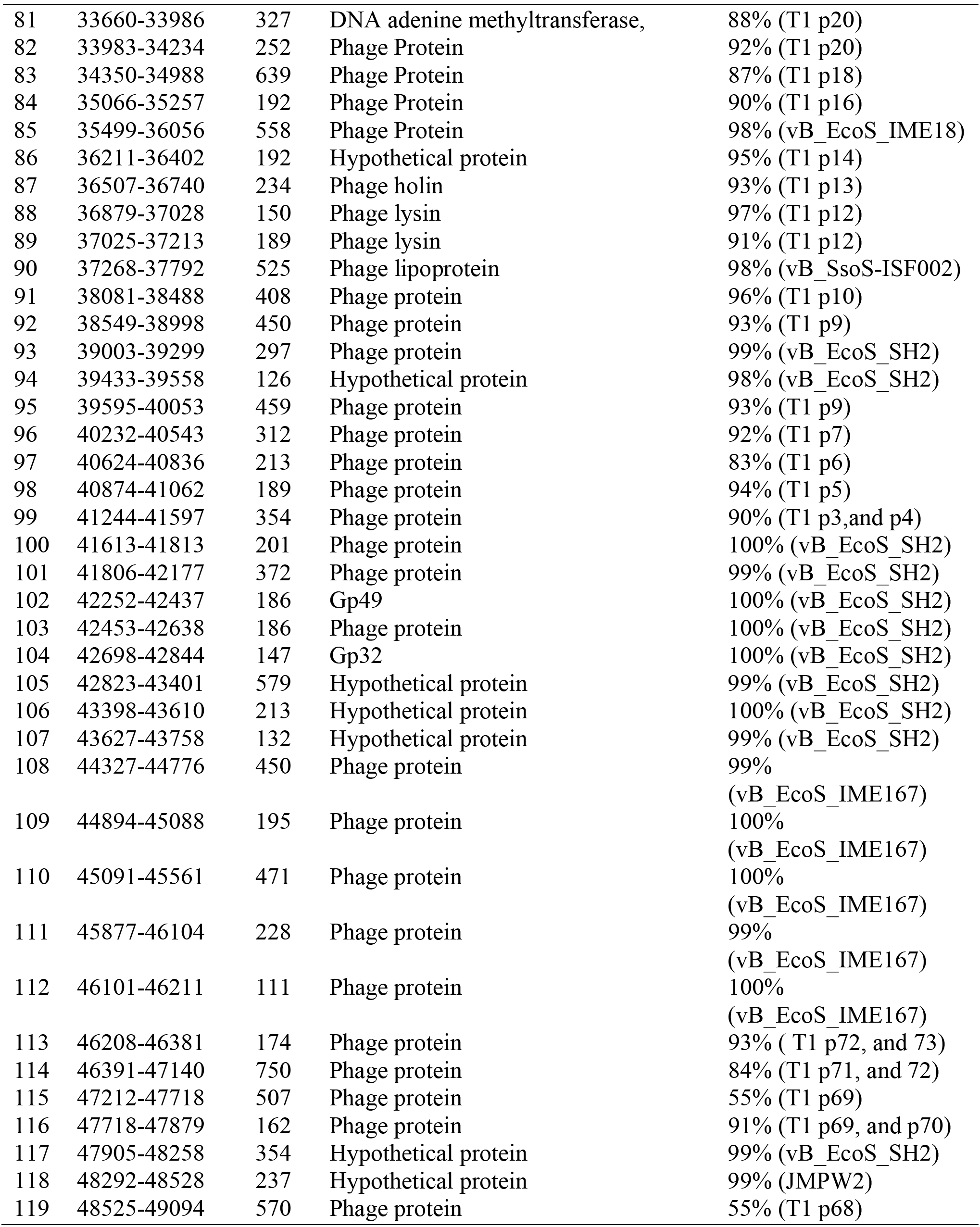
Lytic phage SW1 genes. See also Supplemental Fig. 5 for the circular map.

**Supplemental Table 4.** Cell numbers in the demarcation line as determined by Live/Dead staining.

|  | **Cell number in**<br>**the demarcation**<br>**line** | **Cell number in**<br>**the Halo** | **Fold change** |
| --- | --- | --- | --- |
| BW25113 (SW1)/pCA24N-yfdM<br>vs. BW25113 (SW1)/pCA24N | 160 ± 80 | 1664 ± 90 | 10.4 ± 0.5 |
| MG1655 vs. BW25113 (SW1) | 23 ± 1 | 148 ± 27 | 6.4 ± 0.2 |

**Supplemental Fig. 1.**
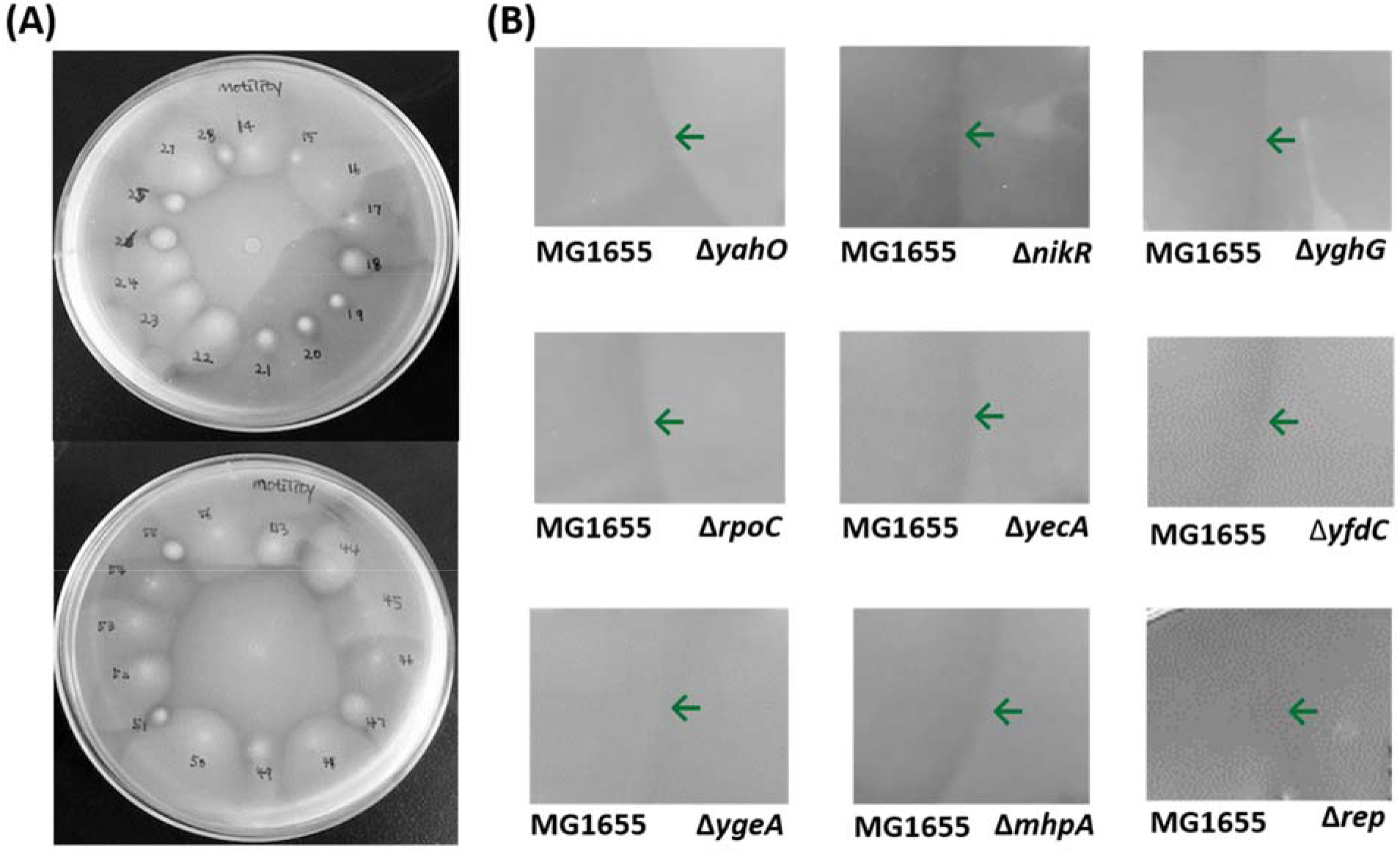
Intersection of motility halos during screening of the Keio collection for proteins involved in kin recognition. **(A)** Screening plates with MG1655 inoculated at the center of the motility plate and members of the Keio collection inoculated on the edge of the plates. **(B)** Re-screening for the formation of a demarcation line by assaying the intersection of the motility halos for MG1655 ^36^ and an individual Keio collection mutant on the right. Each plate contains two strains and representative images are shown. The green arrow indicates where the two swimming halos merge (no demarcation line).

**Supplemental Fig. 2.**
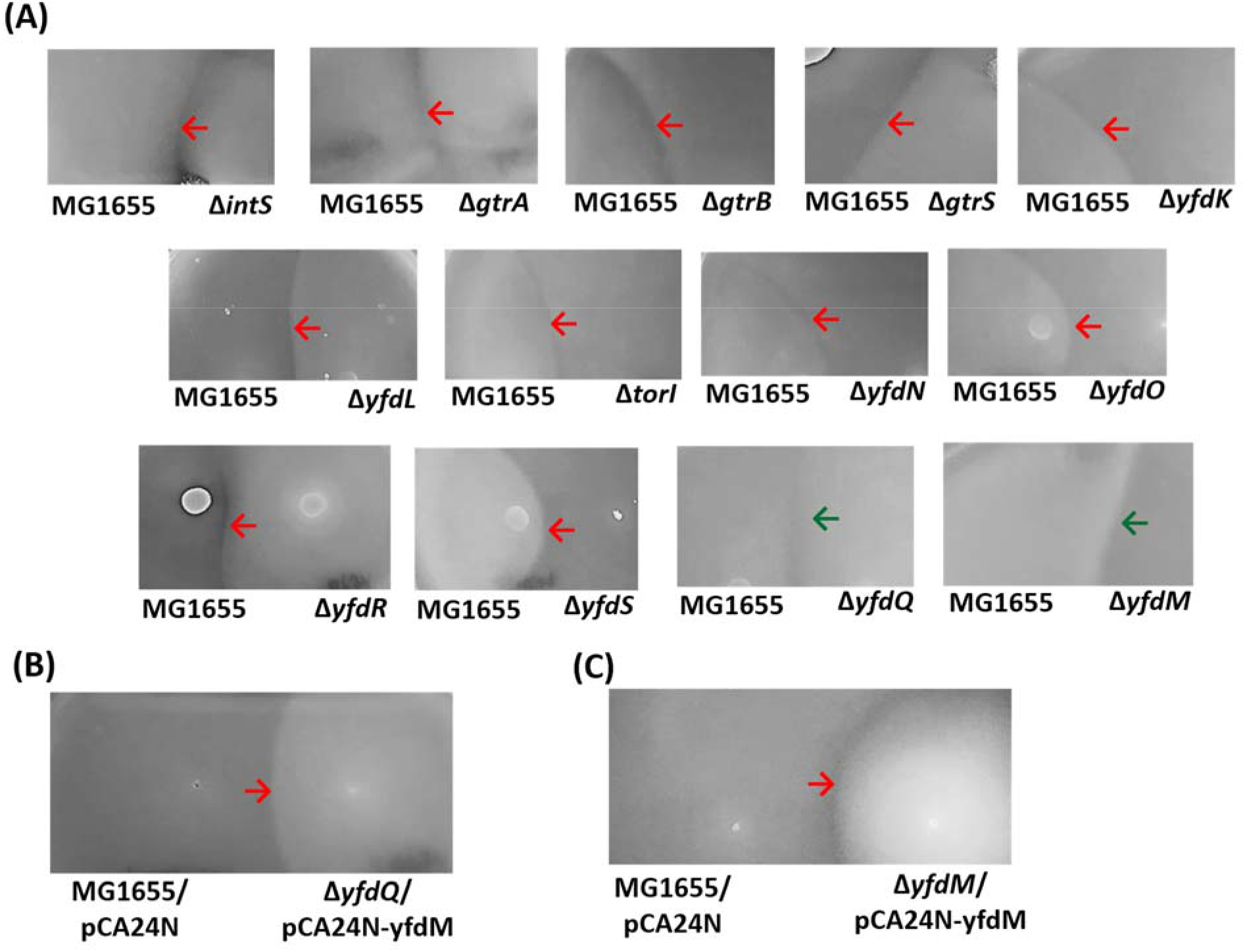
Intersection of motility halos during screening of CPS-53 proteins involved in kin recognition. **(A)** Thirteen single gene mutants of CPS-53 (right) were assayed for the formation of the demarcation line with MG1655. Intersection of the swimming motility halos of **(B)** MG1655/pCA24N vs. Δ*yfdQ*/pCA24N-yfdQ and **(C)** MG1655/pCA24N vs. Δ*yfdM*/pCA24N-yfdM. YfdQ and YfdM were produced from pCA24N-yfdQ and pCA24N-yfdM, respectively, via induction with 0.5 mM IPTG. Each plate contains two strains and representative images are shown. The green arrow indicates where the two swimming halos merge while the red arrow indicates the demarcation line.

**Supplemental Fig. 3.**
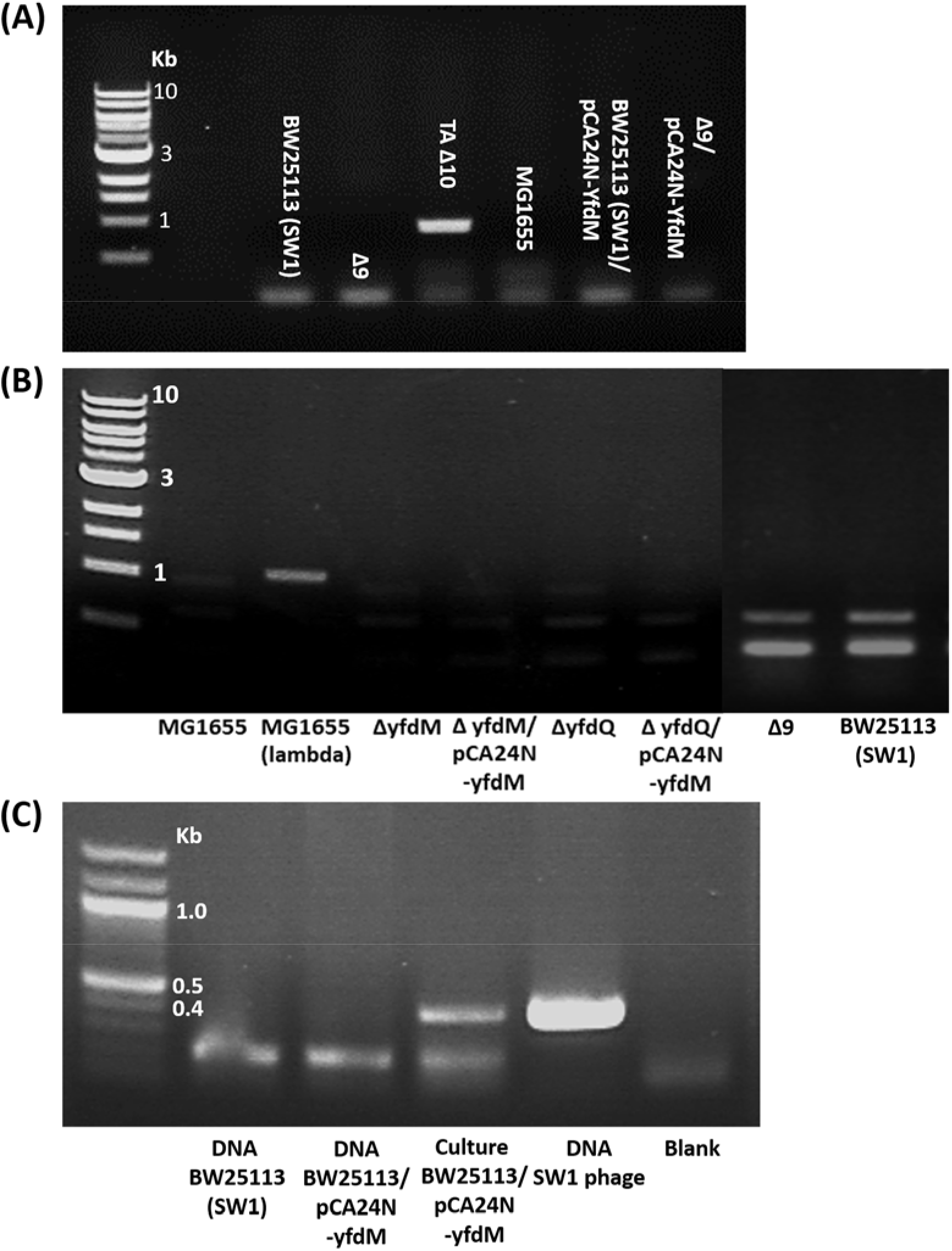
PCR assay for phage ϕ80 and lambda. **(A)** PCR gene fragments to detect for the presence of ϕ80 prophage in the *E. coli* K-12 chromosome. The reaction was performed with primers prAH1500 and prAH1521 for *pinL-tonB*^42^ (**Supplemental Table S2**). TA Δ10 served as the positive control for ϕ80. **(B)** PCR gene fragments to detect the presence of lambda prophage in the *E. coli* K-12 chromosome. The reaction was performed with primers Lambda 1 and Lambda 2 (**Supplemental Table S2**). MG1655 with lambda served as the positive control. **(C)** PCR gene fragments to detect the presence of SW1 phage in the *E. coli* BW25113 chromosome. The reaction was performed with primer recombinase F and recombinase R (Supplemental Table S2). Positive reactions were seen with isolated SW1 DNA and cultures with SW1 YfdM overproduced (BW25113/pCA24N-yfdM); negative reactions occurred for genomic DNA isolated from BW25113(SW1) and BW25113/pCA24N-yfdM.

**Supplemental Fig. 4.**
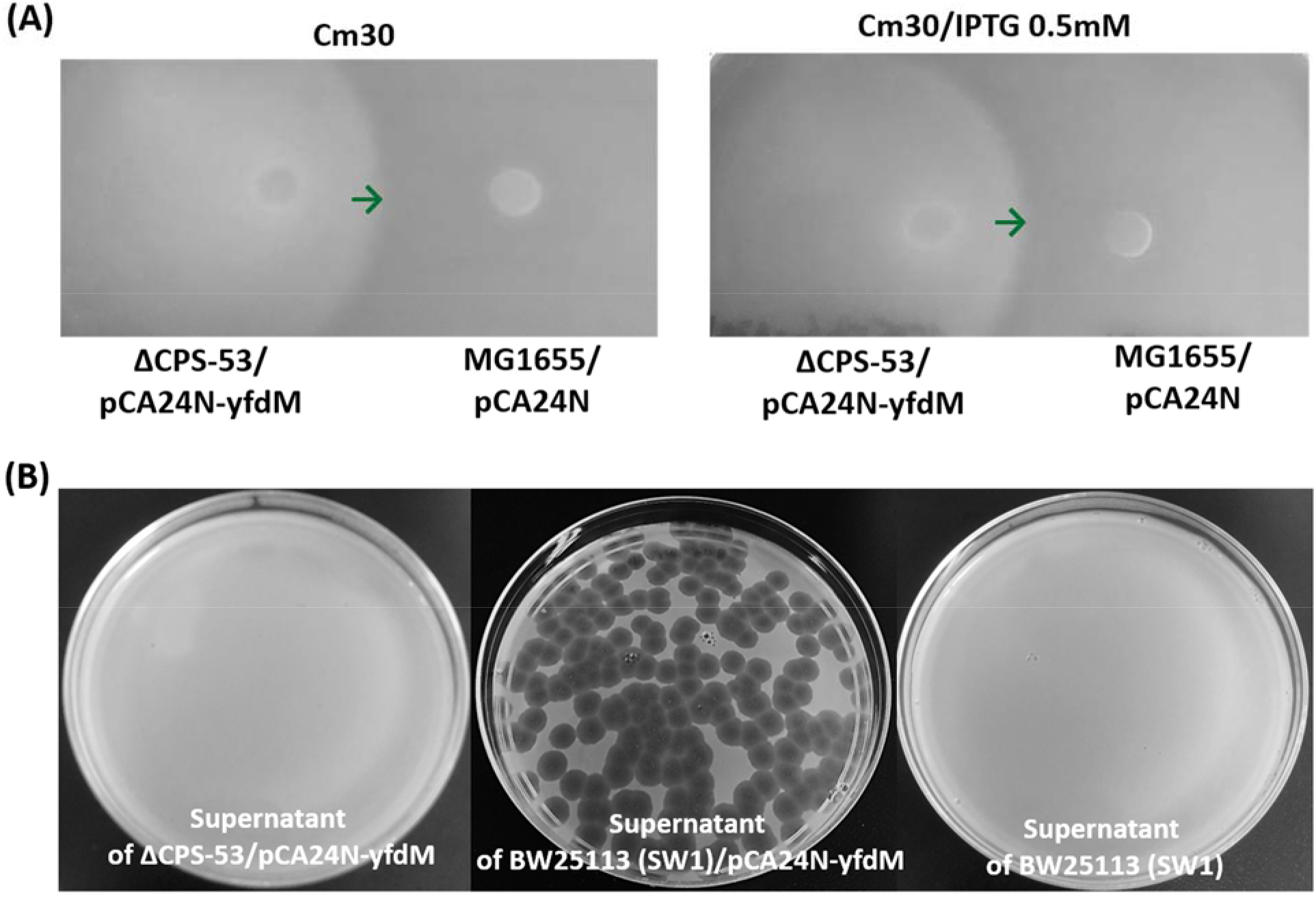
Production of YfdM in the absence of CPS-53 eliminates the demarcation line with MG1655. **(A)** Intersection of the swimming motility halos of ΔCPS-53/pCA24N-yfdM vs. MG1655/pCA24N. Left: no IPTG and right: 0.5 mM IPTG for inducing YfdM production from pCA24N-yfdM. Each plate contains two strains and representative images are shown. The green arrow indicates where the two swimming halos merge (no demarcation line). **(B)** Plaque assay with BW25113 (SW1) cells in soft agar for the supernatant of ∆CPS-53/pCA24N-yfdM, for the supernatant of BW25113/pCA24N-yfdM (middle), and for the supernatant of BW25113 (SW1) (right). Phage stock (100 μL supernatant) and 0.4 mL of BW25113 (SW1) (10^8^ CFU/ml) were added to 4 mL soft agar which was poured onto the LB agar plate. Representative images are shown.

**Supplemental Fig. 5.**
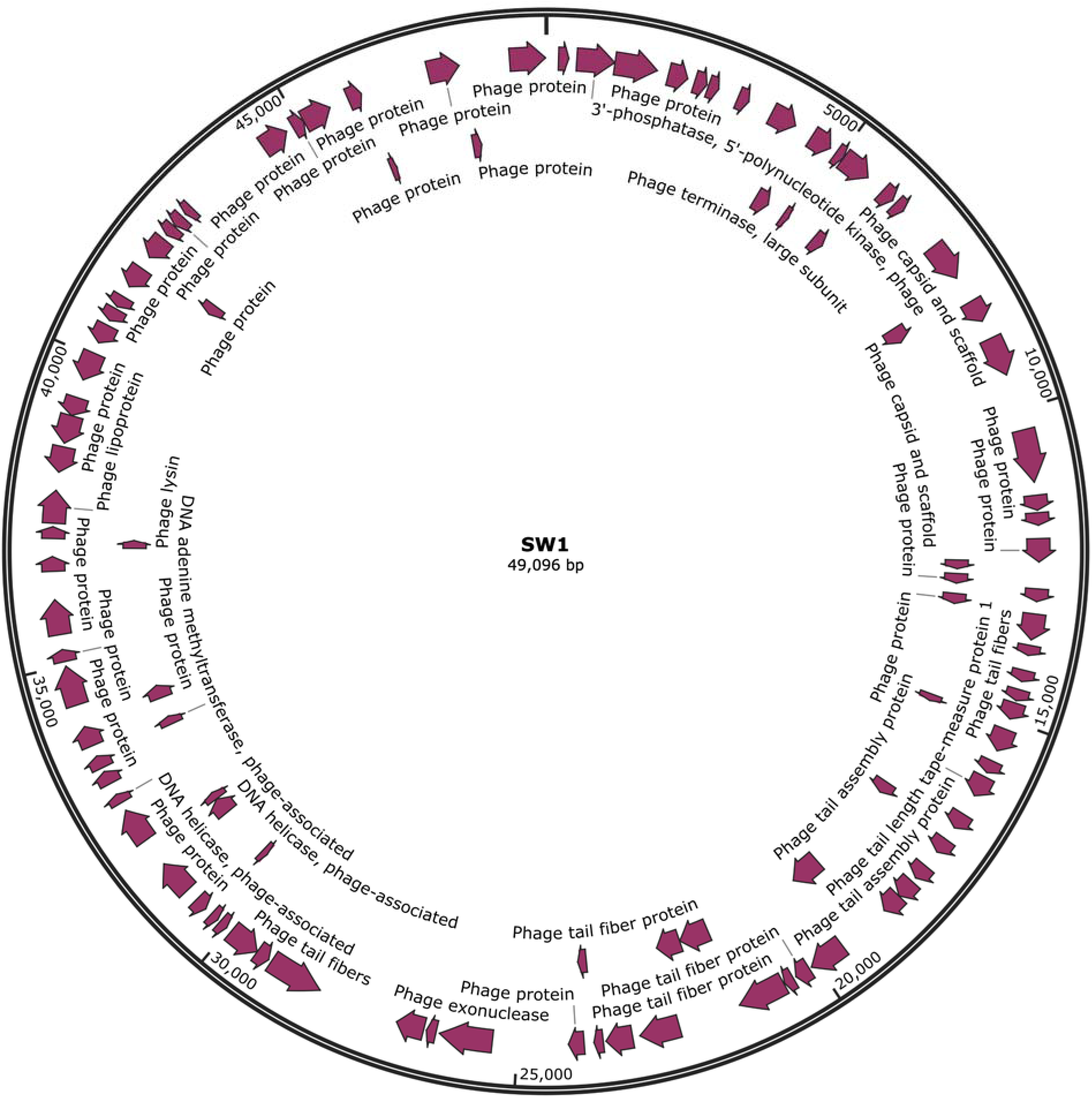
Circular map of the SW1 phage genome. The predicted function of some coding sequences is indicated (see Supplemental Table 3 for more details on individual genes).

**Supplemental Fig. 6.**
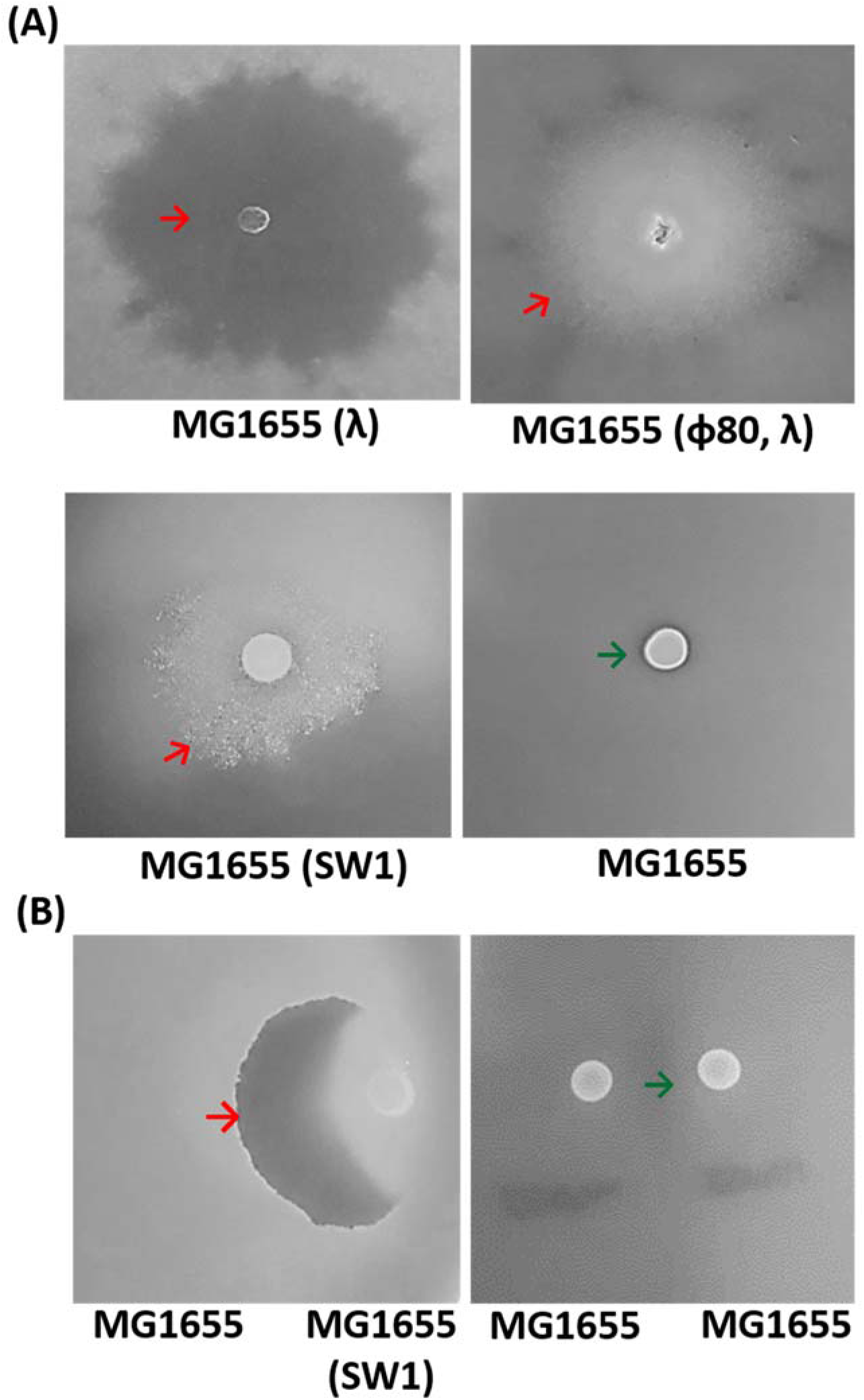
Production of a colony demarcation line for MG1655 colonies harboring phage λ, φ80, and SW1 and conversion of MG1655 into non-kin via SW1. (**A**) Clearance area surrounding colonies on motility plates (colony demarcation line) formed for MG1655 (λ), MG1655(λ + φ80), and MG1655 (SW1), while there is no colony demarcation line for MG1655 that lacks phage. (**B**) Intersection of swimming motility halos of MG1655 (SW1) vs. MG1655 and MG1655 vs. MG1655. Representative images are shown, and motility plates were incubated for 16 h.

**Supplemental Fig. 7.**
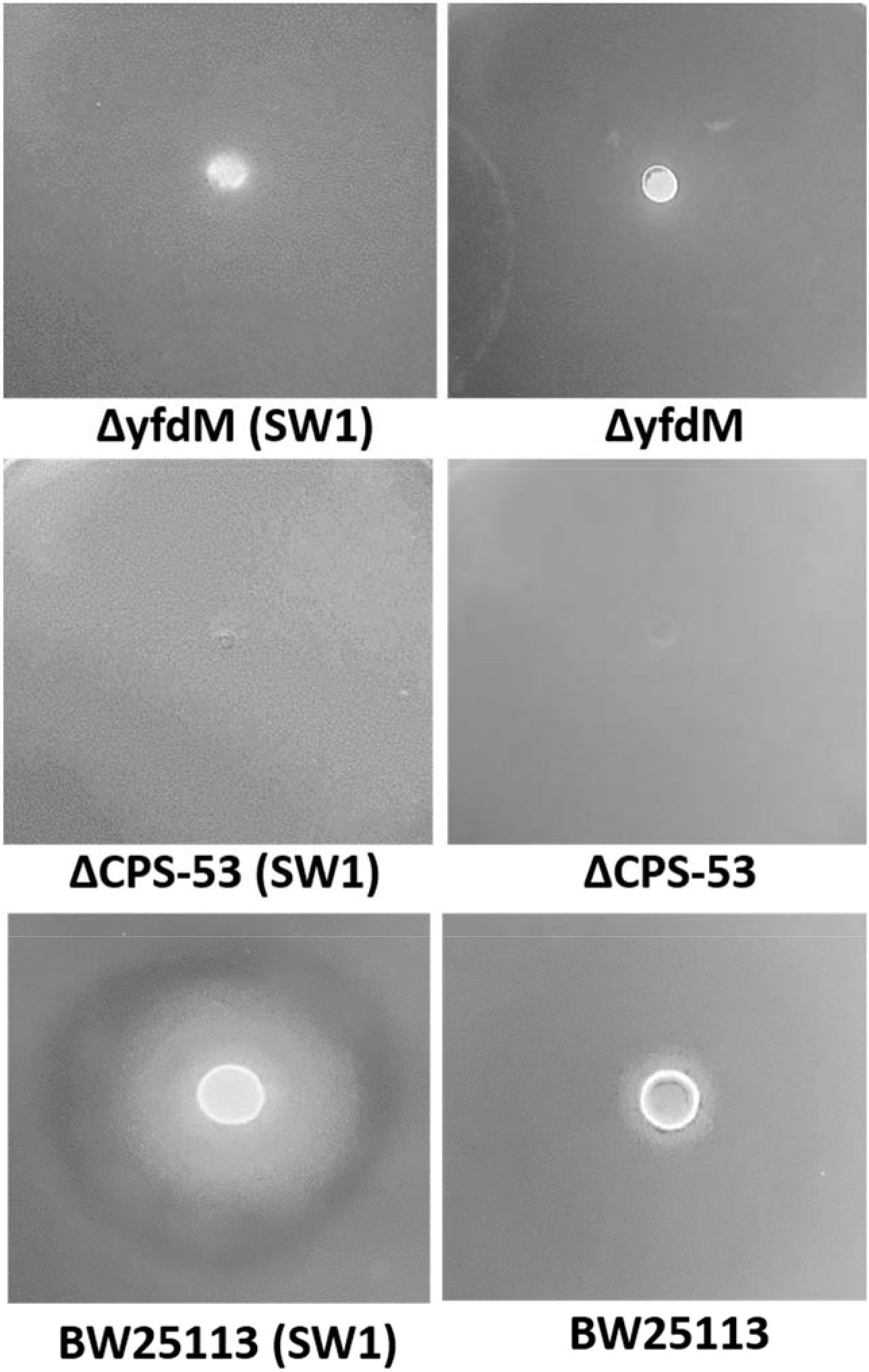
No colony demarcation line by SW1 phage in the absence of CPS-53 and YfdM. Absence of the colony demarcation line on motility plates for BW25113 strains infected with SW1 (indicated by “(SW1)”. Strains lacking SW1 are shown for comparison. Representative images are shown, and the motility plates were incubated for 16 h.

**Supplemental Fig. 8.**
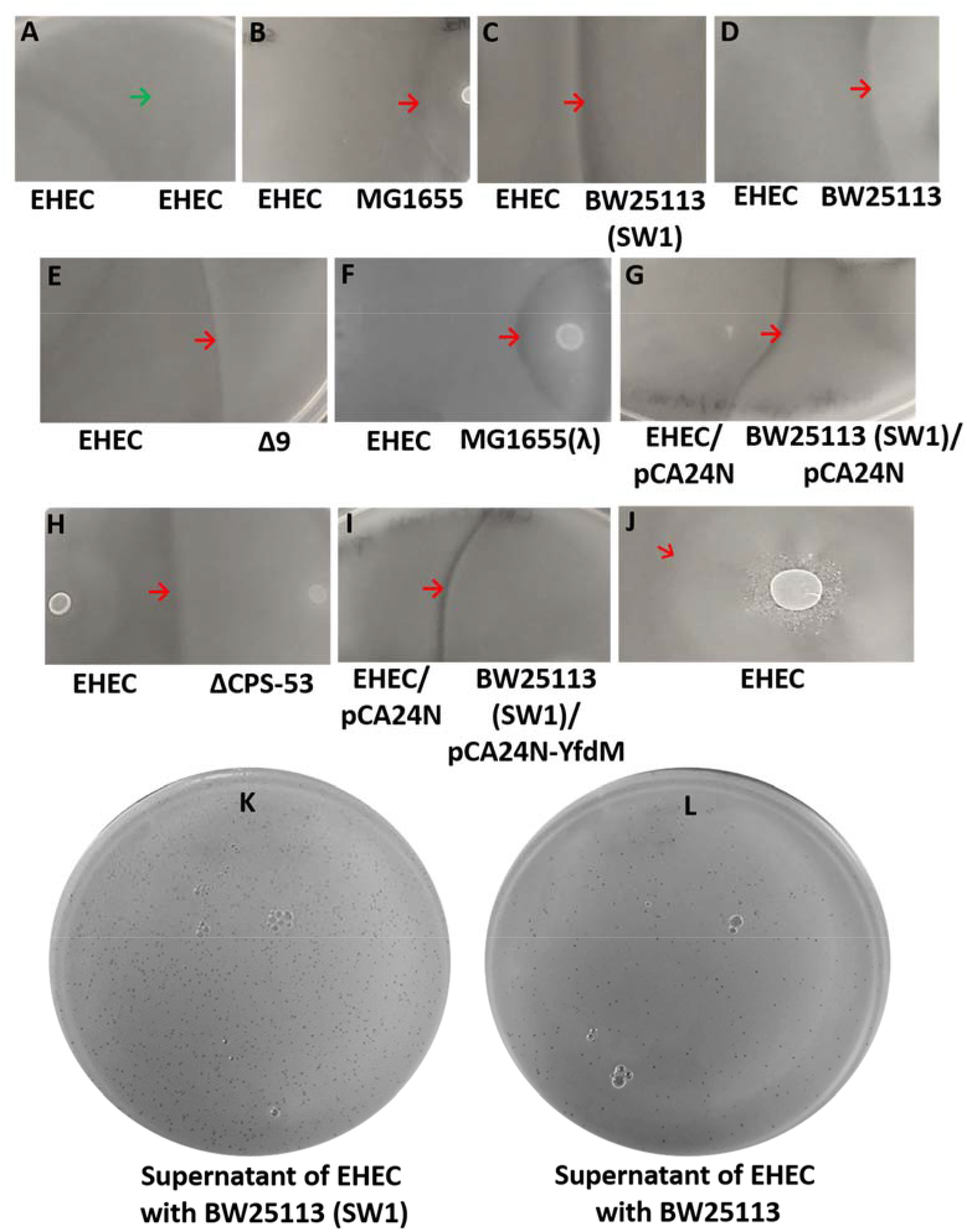
Demarcation lines of EHEC develop between non-kin for swimming cells. Merging of two swimming motility halos for kin (**A**) EHEC vs. EHEC. Demarcation line from the intersection of two swimming motility halos for non-kin (**B**) EHEC vs. MG1655, (**C**) EHEC vs. BW25113 (SW1), (**D**) EHEC vs. BW25113 (**E**) EHEC vs. deleting all the cryptic prophage (BW15113 Δ9), (**F**) EHEC vs. MG1655 with phage lambda, (**G**) EHEC/pCA24N vs. BW25113 (SW1)/pCA24N (**H**) EHEC vs. ∆CPS-53, (**I**) EHEC/pCA24N vs. BW25113 (SW1)/pCA24N-YfdM. Each plate contains two strains. (**J**) Motility halo of an EHEC colony. The green arrow indicates where the two swimming halos merge while the red arrow indicates the demarcation line (**B** through **I**) or colony-demarcation line (**J**). Plaque assay with BW25113 (SW1) cells **(K)** and BW25113 cells **(L)** in soft agar for the supernatant of EHEC. EHEC phage stock and 0.4 mL of BW251113 (SW1) and BW25113 (10^8^ CFU/ml) were added to 4 mL soft agar which was poured onto the LB agar plate. Representative images are shown.

**Supplemental Fig. 9.**
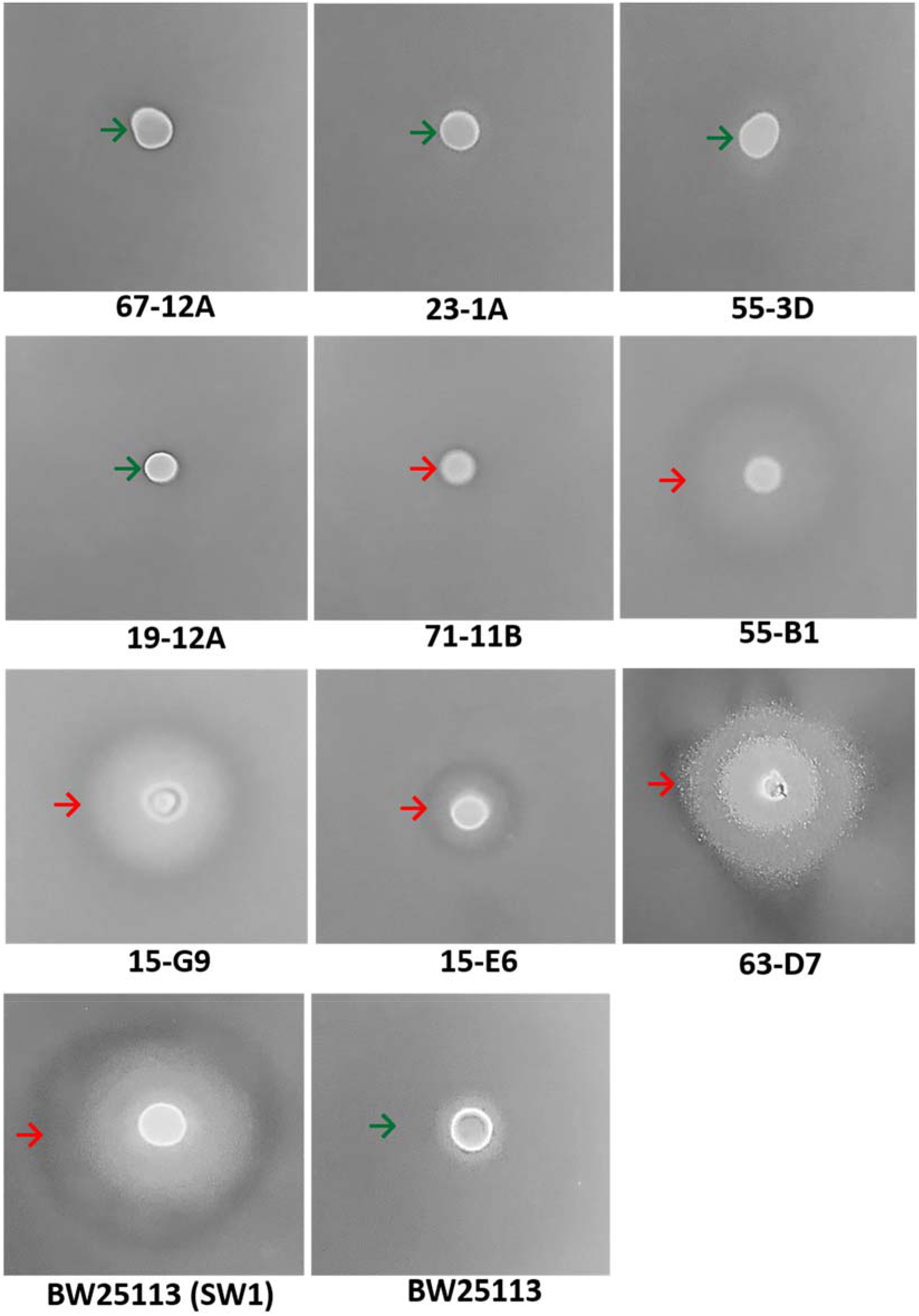
The origin of the SW1 phage is the Keio library. Motility assay of nine random Keio mutants (plate number and well number indicated for each). The green arrow indicates the lack of cell lysis while the red arrow indicates the cell lysis (evidence of phage SW1). Representative images are shown and motility plates were incubated for 16 h.

## References

1. Armitage, J.P. Bacterial motility and chemotaxis. Sci. Progress 76, 451–457 (1992).

2. Wood, T.K., Barrios, A.F.G., Herzberg, M. & Lee, J. Motility influences biofilm architecture in *Escherichia coli*. Appl. Microbiol. Biotechnol. 72, 361–367 (2006).

3. Petrova, O.E. & Sauer, K. Escaping the biofilm in more than one way: desorption, detachment or dispersion. Curr. Opin. Microbiol. 30, 67–78 (2016).

4. Alberti, L. & Harshey, R.M. Differentiation of *Serratia marcescens* 274 into swimmer and swarmer cells. J. Bacteriol. 172, 4322–4328 (1990).

5. Strassmann, J.E., Gilbert, O.M. & Queller, D.C. Kin Discrimination and Cooperation in Microbes. Annu. Rev. Microbiol. 65, 349–367 (2011).

6. Wall, D. Kin Recognition in Bacteria. Annu. Rev. Microbiol. 70, 143–160 (2016).

7. Troselj, V., Cao, P. & Wall, D. Cell-cell recognition and social networking in bacteria. Environ. Microbiol. 20, 923–933 (2018).

8. Gibbs, K.A., Urbanowski, M.L. & Greenberg, E.P. Genetic Determinants of Self Identity and Social Recognition in Bacteria. Science 321, 256–259 (2008).

9. Cardarelli, L., Saak, C. & Gibbs, K.A. Two Proteins Form a Heteromeric Bacterial Self-Recognition Complex in Which Variable Subdomains Determine Allele-Restricted Binding. mBio 6, e00251–00215 (2015).

10. Lyons, Nicholas A., Kraigher, B., Stefanic, P., Mandic-Mulec, I. & Kolter, R. A Combinatorial Kin Discrimination System in *Bacillus subtilis*. Curr. Microbiol. 26, 733–742 (2016).

11. Dey, A. et al. Sibling Rivalry in *Myxococcus xanthus* Is Mediated by Kin Recognition and a Polyploid Prophage. J. Bacteriol. 198, 994–1004 (2016).

12. Wang, X. et al. Cryptic prophages help bacteria cope with adverse environments. Nat. Commun. 1, 147 (2010).

13. Domka, J., Lee, J., Bansal, T. & Wood, T.K. Temporal gene-expression in *Escherichia coli* K-12 biofilms. Environ. Microbiol. 9, 332–346 (2007).

14. Wang, X., Kim, Y. & Wood, T.K. Control and Benefits of CP4-57 Prophage Excision in *Escherichia coli* Biofilms. ISME J 3, 1164–1179 (2009).

15. García Contreras, R., Zhang, X.-S., Kim, Y. & Wood, T.K. Protein Translation and Cell Death: The Role of Rare tRNAs in Biofilm Formation and in Activating Dormant Phage Killer Genes. PLoS ONE 3, e2394 (2008).

16. Webb, J.S., Lau, M. & Kjelleberg, S. Bacteriophage and phenotypic variation in *Pseudomonas aeruginosa* biofilm development. J. Bacteriol. 186, 8066–8073 (2004).

17. Baba, T. et al. Construction of *Escherichia coli* K-12 in-frame, single-gene knockout mutants: the Keio collection. Mol. Syst. Biol. 2, 2006.0008 (2006).

18. Bäumler, A.J. & Sperandio, V. Interactions between the microbiota and pathogenic bacteria in the gut. Nature 535, 85 (2016).

19. Girgis, H.S., Liu, Y., Ryu, W.S. & Tavazoie, S. A Comprehensive Genetic Characterization of Bacterial Motility. PLoS Genet. 3, e154 (2007).

20. Calendar, R., Lindqvist, B., Sironi, G. & Clark, A.J. Characterization of REP− mutants and their interaction with P2 phage. Virology 40, 72–83 (1970).

21. Asadulghani, M. et al. The Defective Prophage Pool of *Escherichia coli* O157: Prophage–Prophage Interactions Potentiate Horizontal Transfer of Virulence Determinants. PLOS Pathog. 5, e1000408 (2009).

22. Langenscheid, J., Killmann, H. & Braun, V. A FhuA mutant of Escherichia coli is infected by phage T1-independent of TonB. FEMS Microbiol. Lett. 234, 133–137 (2004).

23. Böhm, J. et al. FhuA-mediated phage genome transfer into liposomes: a cryo-electron tomography study. Curr. Biol. 11, 1168–1175 (2001).

24. Ogura, Y. et al. The Shiga toxin 2 production level in enterohemorrhagic *Escherichia coli* O157:H7 is correlated with the subtypes of toxin-encoding phage. Sci. Rep. 5, 16663 (2015).

25. Sternberg, N. & Coulby, J. Cleavage of the bacteriophage P1 packaging site (pac) is regulated by adenine methylation. Proc. Natl. Acad. Sci. 87, 8070–8074 (1990).

26. Fujisawa, H. & Morita, M. Phage DNA packaging. Genes to Cells 2, 537–545 (1997).

27. Roberts, M.D., Martin, N.L. & Kropinski, A.M. The genome and proteome of coliphage T1. Virology 318, 245–266 (2004).

28. Ptashne, M. et al. Autoregulation and function of a repressor in bacteriophage lambda. Science 194, 156–161 (1976).

29. Thieffry, D. & Thomas, R. Dynamical behaviour of biological regulatory networks—II. Immunity control in bacteriophage lambda. Bull. Math. Biol. 57, 277–297 (1995).

30. Conway, T. & Cohen, P.S. Commensal and Pathogenic *Escherichia coli* Metabolism in the Gut. Microbol. Spectr. 3, MBP-0006-2014 (2015).

31. Gorbach, S.L. in Medical Microbiology, Edn. 4th. (ed. S. Baron) (Galveston, TX; 1996).

32. Breitbart, M. et al. Metagenomic Analyses of an Uncultured Viral Community from Human Feces. J. Bacteriol. 185, 6220–6223 (2003).

33. Deininger, P. (Academic Press, 1990).

34. Eisenstark, A. Bacteriophage Techniques. Methods in Virology 1, 449–524 (1967).

35. Laver, T. et al. Assessing the performance of the Oxford Nanopore Technologies MinION. Biomolecular Detection and Quantification 3, 1–8 (2015).

36. Kim, W. et al. Identification of an Antimicrobial Agent Effective against Methicillin-Resistant *Staphylococcus aureus* Persisters Using a Fluorescence-Based Screening Strategy. PLoS ONE 10, e0127640 (2015).

37. Maisonneuve, E., Shakespeare, L.J., Jørgensen, M.G. & Gerdes, K. Bacterial persistence by RNA endonucleases. Proc. Natl. Acad. Sci. 108, 13206–13211 (2011).

38. Strockbine, N.A. et al. Two toxin-converting phages from *Escherichia coli* O157:H7 strain 933 encode antigenically distinct toxins with similar biologic activities. Infect. Immun. 53, 135–140 (1986).

39. Kitagawa, M. et al. Complete set of ORF clones of *Escherichia coli* ASKA library (A Complete Set of E. coli K-12 ORF Archive): Unique Resources for Biological Research. DNA Res. 12, 291–299 (2005).

40. Hansen, M.C., Palmer, R.J., Jr, Udsen, C., White, D.C. & Molin, S. Assessment of GFP fluorescence in cells of *Streptococcus gordonii* under conditions of low pH and low oxygen concentration. Microbiology 147, 1383–1391 (2001).

41. Hong, S.H. et al. Synthetic quorum-sensing circuit to control consortial biofilm formation and dispersal in a microfluidic device. Nat. Commun. 3, 613 (2012).

42. Harms, A., Fino, C., Sørensen, M.A., Semsey, S. & Gerdes, K. Prophages and Growth Dynamics Confound Experimental Results with Antibiotic-Tolerant Persister Cells. mBio 8, e01964–01917 (2017).

